# Subtyping psychotic disorders using a data-driven approach reveals divergent cortical and cellular signatures

**DOI:** 10.1101/2025.08.26.672411

**Authors:** Lauren D. Hill, Xi-Han Zhang, Baxter P. Rogers, Anna S. Huang, Victoria Fox, Brandee Feola, Stephan Heckers, Avram J. Holmes, Neil D. Woodward

## Abstract

Structural brain abnormalities in psychosis are well-replicated but heterogenous posing a barrier to uncovering the pathophysiology, etiology, and treatment of psychosis. To parse neurostructural heterogeneity and assess for the presence of anatomically-derived subtypes, we applied a data-driven method, similarity network fusion (SNF), to structural neuroimaging data in a broad cohort of individuals with psychosis (schizophrenia spectrum disorders (SSD) n=280; bipolar disorder with psychotic features (BD) n=101). SNF identified two transdiagnostic subtypes in psychosis (subtype 1: n=158 SSD, n=75 BD; subtype 2: n=122 SSD, n=26 BD) that exhibited divergent patterns of abnormal cortical surface area and subcortical volumes. Compared to controls (n=243), subtype 1 showed moderate enlargement of surface area in frontal and parietal areas and larger dorsal striatal volumes, whereas subtype 2 demonstrated markedly smaller surface areas in frontal and temporal areas and subcortical volumes, including hippocampus, amygdala, thalamus and ventral striatum. When comparing subtypes on clinical characteristics, subtype 2 had more severe negative symptoms, greater neuropsychological impairment, and lower estimated premorbid intellectual functioning compared to subtype 1. Integrating cell-type data imputed from gene expression in the Allen Human Brain Atlas revealed an association between interregional reductions in surface area and layer 5 glutamatergic neuron abundance, critical for corticostriatal network connectivity and cognitive function, whereas reductions in cortical thickness spatially coupled with glia cell and interneuron abundance, in subtype 2. These outcomes indicate that regional variations in surface area, linked to different cell-types than cortical thickness, may be an important biomarker for understanding the pathophysiological trajectories of psychotic disorders.

## INTRODUCTION

Psychosis disrupts how an individual thinks, feels, and perceives the world. Approximately 1.5 to 3.5% of the population suffers from a psychotic disorder, with schizophrenia spectrum disorders (SSD) and bipolar disorder with psychotic features (BD) being the most severe [1]. Over four decades of neuroimaging research has revealed numerous brain abnormalities in people with psychosis providing critical insights into the pathophysiology, etiology, and treatment of these disorders [2–4]. Alterations in brain structure are among the most prominent findings and include lower cortical thickness in frontal and temporal areas; and smaller subcortical volumes, especially the hippocampus and thalamus [5]. Abnormalities in brain structure are well-replicated but heterogeneous, as demonstrated by three observations. First, the effect sizes are relatively small, ranging from approximately 0.20 to 0.40 standard deviations below normal, indicating significant overlap with healthy individuals [6, 7]. Second, the volumes of many brain structures are more variable in psychotic disorders compared to healthy individuals [8, 9]. Third, recent studies using normative modelling have revealed that while the majority of patients have at least one brain region that falls outside the normal range, only up to approximately 30% of patients demonstrate abnormal deviations for any given brain structure [10, 11]. Combined, these findings strongly suggest the presence of subtypes of psychosis characterized by different patterns of brain abnormalities and may provide clues for distinct illness mechanisms.

The most common approach to dissecting neurostructural heterogeneity in psychosis is to examine the effect of diagnosis, usually by comparing SSD and BD, and illness stage (i.e. early stage vs. chronic). As SSD and psychotic BD have both shared and distinct clinical symptoms, it is expected that biological similarities and differences can be found. Broadly, structural brain abnormalities are often quantitatively larger in SSD, but qualitatively similar across disorders. For example, both SSD and BD are associated with fronto-temporo-occipital grey matter deficits, however, in SSD, these abnormalities are more severe in a number of regions including the insula [12, 13]. The modest accuracy of machine learning methods at discriminating between SSD and BD based on brain features underscores the similarities between disorders [14]. A similar pattern emerges when comparing early and chronic stages of psychosis. While, the magnitude of the abnormalities is often larger in chronic patients, there is often considerable overlap between early and chronic stages of the illness [15–17]. Consequently, diagnostic and illness stage appear to account for a modest portion of the variance in brain structure abnormalities in psychosis.

To make progress, investigators are turning towards data-driven approaches to parsing heterogeneity. Studies have examined brain structure in subgroups defined based on relatively low dimensional clinical, cognitive, and neurophysiological data [18–20]. The Bipolar and Schizophrenia Network for Intermediate Phenotypes (BSNIP) consortium, which identified three ‘biotypes’ in a large sample of individuals with SSD and BD based on a combination of cognitive and neurophysiological biomarkers, is perhaps the most prominent example. Biotypes 1 and 2, which demonstrated more severe impairments on biomarkers of neurophysiology and cognition, were associated with widespread reductions in grey matter volume. In contrast, biotype 3, which demonstrated near normal or normal performance on biomarkers of neurophysiology and cognition, exhibited relatively small alterations in grey matter volume [20]. Few studies have applied data-driven approaches to high-dimensional structural neuroimaging data; however, there is growing evidence that there are at least two SSD subtypes [8, 21, 22]. For instance, using a method called HYDRA applied to voxel-wise structural images, Chand et al. [21] identified two subgroups in a cohort of SSD patients; one characterized by widespread lower grey matter volume across cortical and subcortical areas and a second defined by larger basal ganglia volumes in the context of otherwise mostly normal brain structure. The same group subsequently replicated the two subtypes in an independent sample of SSD and linked expression of the subtype associated with widespread smaller grey matter volumes to lower cognitive functioning [21, 22].

Existing studies support the presence of neurostructural subtypes but also have limitations. None included other psychotic disorders such as BD. As noted above, similarities in structural brain abnormalities observed in SSD and BD, along with overlapping symptoms and cognitive deficits, suggests there may be transdiagnostic neurostructural subtypes. Similarly, most focused on chronic patients, or did not report the proportions of individuals in the early stage and chronic stages of illness. Again, qualitative similarities between early stage and chronic samples raises the possibility that neurostructural subtypes may transcend illness stage. Another limitation of the existing literature is that they focused on volume or cortical thickness exclusively. In the case of the cortex, volume represents the product of two dimensions, surface area and thickness. Cortical thickness and surface area have different genetic architectures, exhibit distinct developmental trajectories, and are differentially related to clinically-relevant variables including cognition and educational attainment [23–25]. Regional surface area exhibits higher heritability compared to cortical thickness and is associated with genes critical to early growth and development of the brain, including those that regulate progenitor cell types present in fetal development [26]. Regional cortical thickness is rather associated with genes implicated in cell differentiation, migration, adhesion, and myelination, and may reflect environmental influences [26]. Consequently, focusing on volume exclusively may conflate the contributions of surface area and thickness and therefore not capture the more basic structural elements of the cortex that may aid in the understanding of disease mechanisms in psychotic disorders.

The current study sought to replicate and build on prior studies in several ways. First, we included a relatively large cohort of individuals with psychosis spanning the early and chronic stages of both SSD and BD. This allows us to simultaneously determine if there are subtypes that transcend diagnostic boundaries and illness stage and increase generalization to the broader population of psychosis. Second, we used a relatively new data-driven method, similarity network fusion (SNF) [27], to identify subtypes of psychosis individuals based on multiple features of brain structure, including cortical thickness and surface area, and subcortical volumes. SNF is designed to specifically capture both common and complementary information from different data types providing insight into how informative each feature is to the observed subtypes. Third, the rich phenotypic data available on our psychosis cohort allows us to characterize out-of-model features including clinical symptoms, neuropsychological functioning, and illness antecedents such as premorbid intellectual and psychosocial functioning. Finally, we leveraged microarray data from the Allen Human Brain Atlas and single-nucleus RNA-sequencing data to examine the cell-type correlates of large-scale morphometric pathology seen in subtypes of psychosis and bridge gaps across levels of biological systems.

We hypothesized that we would replicate prior studies by identifying two or more anatomically-derived subtypes with at least one demonstrating relatively large and widespread abnormalities in brain structure. We further sought to extend prior data-driven subtyping studies by: 1) establishing which brain features, cortical thickness, surface area, subcortical volumes, or a combination of these features, distinguish subgroups; 2) determining the extent to which anatomically-derived subtypes align with diagnosis and illness stage; 3) clarifying the clinical phenotypes associated with brain-based subtypes; and 4) establishing whether interregional profiles of psychosis subtype neuroanatomical abnormalities spatially relate to cell-type abundances across the brain.

## METHODS

### Participants and study procedures

Data for the current study was drawn from a repository containing 756 individuals that participated in one of three neuroimaging studies (CT00762866; R01MH070560; R01MH102266) conducted in the Department of Psychiatry and Behavioral Sciences at Vanderbilt University Medical Center (VUMC). All study procedures were approved by the Vanderbilt Institutional Review Board. Each participant provided written informed consent to participate in the separate studies and contribute their data to the data repository. See supplemental material for inclusion and exclusion criteria. Briefly, participants with psychotic disorders (i.e. SSD, BD) and healthy individuals were recruited from the Psychotic Disorders Program at VUMC, and surrounding Nashville area via advertisement, respectively. The psychosis cohort included individuals in both early or chronic stages defined as illness duration ≤ 2 or >2 years, respectively. From the initial cohort of 756, 61 were excluded for data quality reasons and 71 for diagnosis not of interest or ineligible (Figure S1), leaving a final sample of 243 healthy individuals and 381 people with a psychotic disorder (SSD n=280, BD n=101) (Table 1, Table S1).

**Table 1.** Sample demographics and clinical characteristics.

|  |  | Healthy Individuals<br>(n=243) | Psychotic Disorder<br>(n=381) | Statistics |  |  |
| --- | --- | --- | --- | --- | --- | --- |
| | | No. (%) | No. (%) | $\chi^2$ | df | p-value |
| **Sex** (female) |  | 91 (37.45) | 137 (35.96) | 0.09 | 1 | 0.77 |
| **Race** |  |  |  | 5.11 | 4 | 0.27 |
|  | White | 169 (69.55) | 252 (66.14) |  |  |  |
|  | Black | 54 (22.22) | 104 (27.30) |  |  |  |
|  | Asian | 11 (4.53) | 8 (2.10) |  |  |  |
|  | American Indian or Alaskan<br>Native | 3 (1.23) | 4 (1.05) |  |  |  |
|  | Other | 6 (2.47) | 13 (3.41) |  |  |  |
| **Ethnicity** (Hispanic) |  | 12 (4.94) | 17 (4.46) | 5.22 | 2 | 0.07 |
| **Handedness** (right) |  | 223 (91.77) | 342 (89.76) | 0.36 | 1 | 0.55 |
| **Diagnosis** (SSD/BD) |  | --- | 280/101 |  |  |  |
| **Illness-Stage** (early/chronic) |  | --- | 236/144 |  |  |  |
| | | M ± SD | M ± SD | t | df | p-value |
| **Age** (years) | | 28.03 ± 9.70 | 28.36 ± 10.76 | -0.39 | 622 | 0.69 |
| **Education** (years) | | 15.36 ± 2.11 | 13.55 ± 2.18 | 9.96 | 590 | <0.001 |
| **Parental Education** (years) | | 14.64 ± 2.29 | 14.70 ± 2.75 | -0.25 | 567 | 0.80 |
| **Estimated Premorbid IQ** | | 108.50 ± 7.57 | 101.99 ± 10.89 | 8.00 | 596 | <0.001 |
| **SCIP** (z-score) | | 0.14 ± 0.63 | -0.92 ± 0.93 | 15.55 | 610 | <0.001 |
| **Age of Illness Onset** (years) | | --- | 21.71 ± 6.36 | --- | --- | --- |
| **Duration of Illness** (months) | | --- | 75.13 ± 117.02 | --- | --- | --- |
| **PANSS Positive Symptoms** | | --- | 17.31 ± 8.30 | --- | --- | --- |
| **PANSS Negative Symptoms** | | --- | 14.61 ± 6.82 | --- | --- | --- |
| **PANSS General Symptoms** | | --- | 30.61 ± 8.96 | --- | --- | --- |
| **Premorbid Adjustment** (total) | | --- | 1.37 ± 0.68 | --- | --- | --- |
| **CPZ Equivalents** | | --- | 379.80 ± 494.64 | --- | --- | --- |

**Note.** SSD, Schizophrenia Spectrum Disorder; BD, Bipolar with Psychotic Features; SCIP, Screen for Cognitive Impairment in Psychiatry; PANSS, Positive and Negative Syndrome Scale; Italics indicate significant *p*-values. Estimated premorbid IQ measured by the WTAR, Wechsler Test of Adult Reading predicted score. Duration of illness was defined as the time at which an individual first met criteria for psychosis (based on extensive interview, review of medical records, and collateral reports) until the date of study enrollment. CPZ, chlorpromazine equivalent. Premorbid adjustment scale available for 217 individuals with psychosis.

#### Diagnostic and clinical assessments

Diagnoses were confirmed, and ruled out in the case of healthy individuals, by clinical interview with the Structured Clinical Interview for DSM (IV/V) Disorders (SCID) conducted by trained study personnel. Symptoms of psychosis were quantified in the psychosis cohort using the Positive and Negative Syndrome Scale (PANSS) [28]. The Screen for Cognitive Impairment in Psychiatry (SCIP; Purdon, 2005) was administered to all study participants to quantify current neuropsychological functioning. The SCIP includes five sub-tests assessing immediate and delayed verbal learning, verbal fluency, working memory, and processing speed. Subtest raw scores were *z*-scored based on published norms [29] and averaged to create a SCIP composite score. Premorbid intellectual and social functioning were assessed using the Wechsler Test of Adult Reading (WTAR) [30] and Premorbid Adjustment Scale (PAS), respectively [31].

#### MRI data acquisition, quality assurance, and processing

A high resolution T1-weighted anatomical scan (voxel size=1mm^3^ isotropic) collected in the axial plane was obtained on every participant on one of two identical 3T Philips MRI scanners equipped with a 32-channel head coil located at Vanderbilt Institute for Imaging Sciences (VUIIS) (detailed in supplement). All imaging data were stored and processed on the VUIIS Center for Computational Imaging XNAT [32, 33]. T1-weighted images were visually inspected for neuroimaging artifacts. A nuisance variable (‘scan type’) was included as a covariate in all analyses to account for minor differences in the T1 acquisition parameters between studies. The entropy-focus criterion (EFC), a quantitative measure of ghosting and blurring induced by head motion [34], was calculated for each T1-weighted image using MRI Quality Control (MRIQC) [35] and included as a covariate in all imaging analyses. Cortical thickness (CT) and surface area (SA) values (68 regions) and subcortical volumes (SV, 14 regions) were derived from T1-weighted structural scans using the Desikan-Killiany (DK) atlas in FreeSurfer (v7.2.0) [36, 37]. FreeSurfer segmentations were visually inspected, and 26 subjects were excluded for segmentation failures.

### Data integration and clustering

#### Similarity network fusion

The *SNFtool* package v.2.3.1 running in R (version 4.4.1) was used to integrate data types (i.e. CT [68 features], SA [68 features], SV [14 features]) and identify subgroups within the psychosis cohort. Prior to SNF data integration, covariates including age, biological sex, scan type, and EFC from MRIQC [35] were regressed out using general linear models for each brain morphology feature. Residuals were used as input for the SNF analysis. SNF has been previously described [27] (http://compbio.cs.toronto.edu/SNF/SNF/Software.html). Briefly, SNF first creates separate similarity networks for each data type and then integrates these networks using a nonlinear combination method to iteratively fuse the networks for each data type into a single, global network, representing the full spectrum of included features. Across iterations, low weight edges (i.e., similarities between participants) in each data type are removed to reduce noise, while high weight edges are added together. The similarity matrices for each of the three data types (CT, SA, and SV) were calculated using Euclidean distance with a nearest neighbors values of 30 and a normalization parameter of 0.6, based on the ranges recommended in the *SNF tool* R package, 10-30 for K (sample size/10) and 0.3-0.8 for alpha [27]. Cluster number was chosen using the *estimateNumberOfClustersGivenGraph* function to calculate eigen-gaps and rotation costs for each K-alpha combination. We tried 126 combinations of K and alpha hyperparameters (K=10-30; alpha=0.3-0.8), consistent with prior applications of SNF to neuroimaging data [38] to determine stability of suggested cluster number. Normalized mutual information (NMI) was used to determine the relative importance of each included brain feature for determining clustering, where higher NMI scores indicate greater contribution to participant similarity (*rankfeaturesbyNMI* function). NMI is a measure of similarity between a clustering outcome defined by all the model features and a clustering outcome defined by any single model feature of interest. Spectral clustering (*spectralClustering* function) was then applied to identify subgroups based on participant similarity matrices determined based on all 150 features across 1000 iterations of resampling 80% of participants. A silhouette plot, derived from the clustering consensus matrix, quantified the similarity between participants within a given subgroup compared to participants in all other subgroups.

#### Cluster stability

The stability of the clusters obtained from SNF were assessed via resampling-based stability testing with 1000 iterations. For each iteration, 80% of the participants were randomly sampled, their brain morphology features integrated using SNF, and clustered into subgroups. Three metrics of cluster stability were calculated for each iteration, 1) the percentage of time each participant clustered with every other participant (when pairs were included in the same resampling iteration); 2) the Adjusted Rand Index [39, 40] which measures the overlap between clusters to determine stability of the data-driven model (range: 0 to 1, 0=completely random cluster overlap, 1=identical clustering across iterations); and 3) the percentage with which each model feature remained among the top 35 features across iterations based on NMI scores.

### Statistical analyses

Continuously distributed and dichotomous variables were analyzing using independent groups *t*-tests/general linear regression and Chi-square analyses, respectively. Age, biological sex, scan type, scan quality (i.e. EFC), and, in the case of subcortical volumes, estimated total intracranial volume (ICV), were included as covariates in the analysis of brain structure dependent variables derived from FreeSurfer. Multiple comparisons were controlled for using the false discovery rate (FDR) (*p*_FDR_<0.05) [41]. Cohen’s *d* effect sizes for each region of interest were computed based on the group contrast *t*-statistics [42].

### Cell-type imputation and analysis

Cell-type density maps were downloaded from Zhang et al. [43] In brief, gene expression signatures of 24 neuronal and nonneuronal cell-types were derived from single-nucleus RNA-sequencing data [44], then, the signatures were used to impute the fractions of each cell-type from the total gene expression estimated from the microarray data covering the cortex [45]. The 24 cell-type fractions of each microarray sample were registered to the closest DK parcel based on the vertex ID of the sample. Parcel-level cell-type fractions were obtained by taking the average of samples registered to the same parcel.

Cell-types’ combinational correlation to interregional neurostructural deviations was estimated via permutational canonical correlation analysis (PermCCA), which examines the linear combination of all the cell-type interregional abundances that maximally correlate with each subtype neurostructural profile (for both CT and SA). The statistical significance of the canonical variates was tested via a permutation method that controls for cortical spatial autocorrelation, where the null distributions of cell-type fractions were generated from the Baum method [46]. Then, FDR was applied to correct for four comparisons. The cell-types’ linear combinational correlation to each neurostructural profile was measured by the cell-type composite score. Each cell-type’s contribution to this correlation was measured by loadings, the correlation between cell-type abundance distribution and neurostructural profile canonical variate.

## RESULTS

### Brain structure in psychosis

To establish if structural abnormalities in our psychosis cohort are representative of the broader psychosis population, we calculated effect sizes (ES) for the psychosis vs. healthy individuals’ contrast for all brain features and compared them qualitatively and quantitatively to the ENIGMA Schizophrenia meta-analyses of brain structure [6, 7]. As shown in Figure 1A, psychosis was associated with widespread cortical thinning, smaller surface areas, and smaller subcortical volumes. Consistent with ENIGMA Schizophrenia, ESs were in the small to medium range (i.e., 0.20 to 0.40). Correlation analyses revealed that ESs obtained in the current sample correlated strongly with those reported in the ENIGMA Schizophrenia meta-analyses (*r*=0.80, *p*<2.2^-16^, Figure 1D) indicating that the overall pattern of structural abnormalities observed in our sample is similar to the broader psychosis population. These findings suggest that disease-related patterns of neurostructural abnormalities in psychotic disorders are highly replicable.

**Figure 1.**
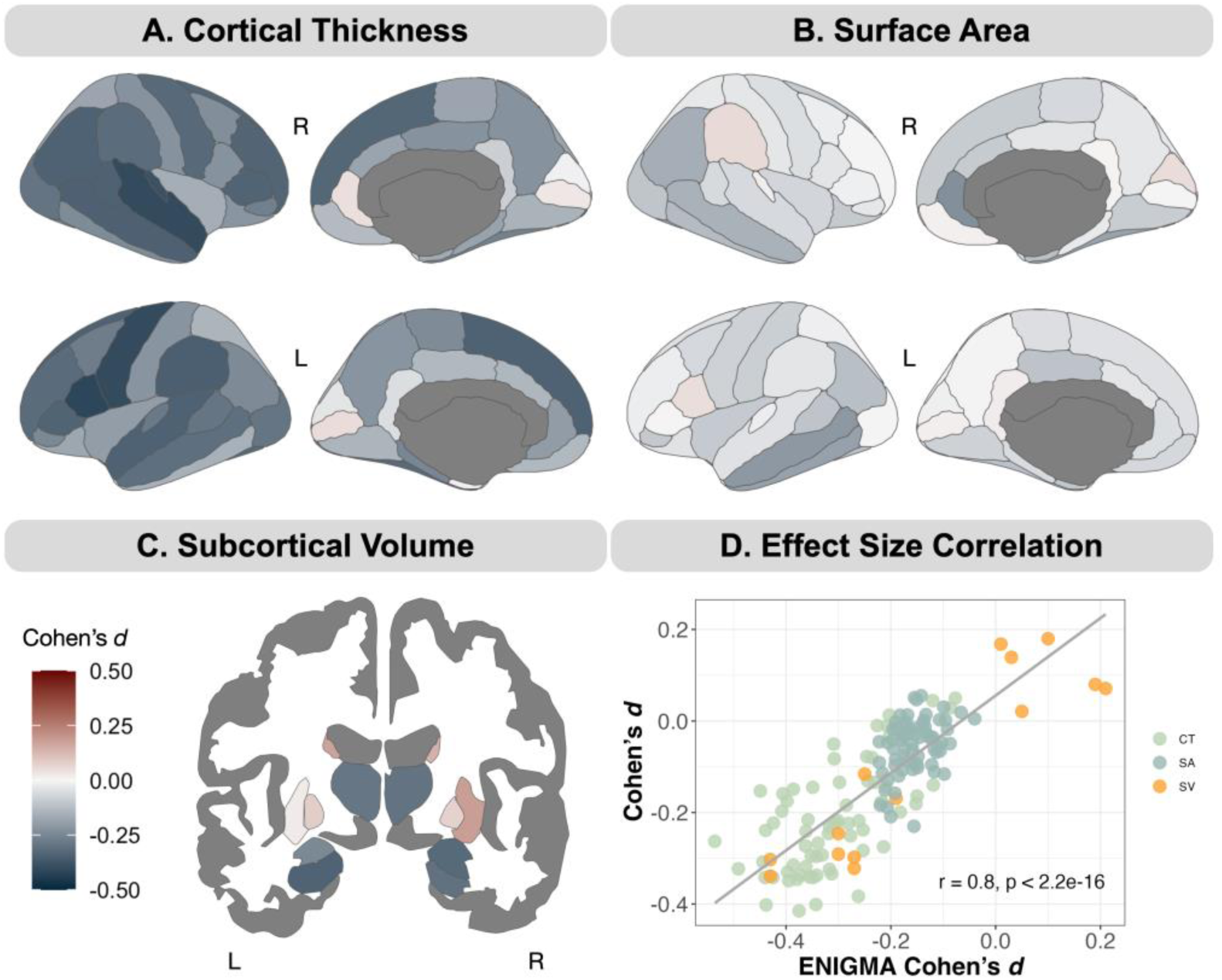
Maps of regional Cohen’s *d* effect sizes for psychosis vs. healthy individual contrast. Panels A-C: regions with lower and higher cortical thickness, cortical surface area, and subcortical volume in psychosis are depicted in cool and warm colors, respectively. Panel D: Pearson’s correlation between regional effect sizes in the current study and ENIGMA Schizophrenia.

### Composition and stability of neuroanatomically-derived clusters in psychosis

SNF and spectral clustering revealed two participant similarity network clusters (Figure 2A) comprised of n=233 and n=148 individuals (hereafter referred to as subtypes 1 and 2, respectively) based on eigen-gaps and rotation costs. The average silhouette width was 0.93 indicating a strong cluster structure (Figure 2A, silhouette plot).

**Figure 2.**
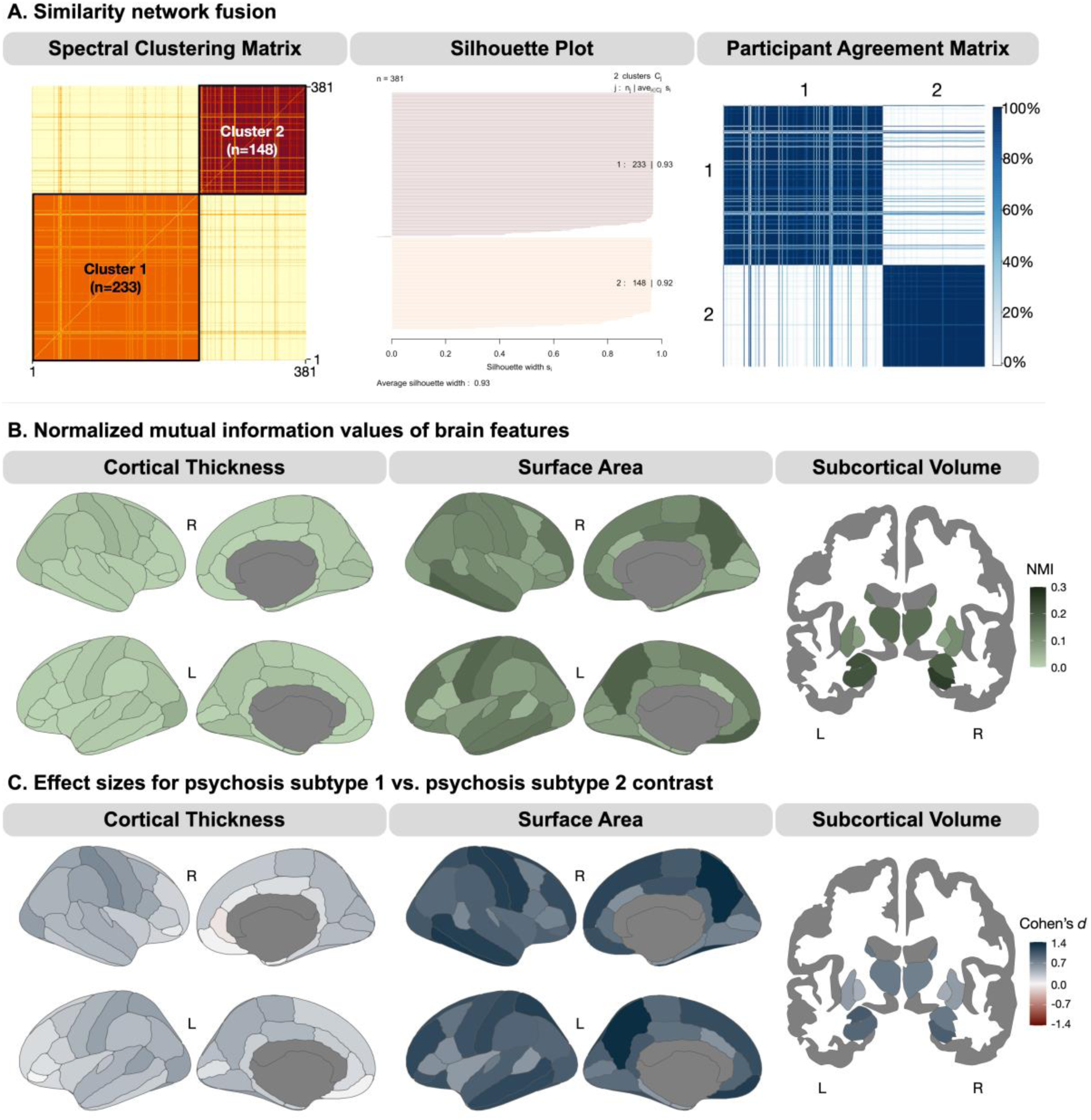
Data-driven similarity network fusion analysis of brain morphology features revealed two clusters in psychosis. Panel A: participant-to-participant SNF-combined similarity matrix after clustering into two subtypes of psychosis individuals. Silhouette plot illustrating the similarities between participants within a cluster versus the other cluster. Cluster number: n of cluster | average cluster silhouette width. Participant agreement matrix, how often each participant clustered with one another across resampling 80% of psychosis individuals 1000x (dark blue = higher percentage of instances that each pair of participants clustered together. Panel B: normalized mutual information (NMI) values across brain regions. Surface area (SA) and subcortical volume (SV) had higher NMI values compared to cortical thickness (CT), indicating greater importance in determining data-driven subtypes. Panel C: the largest Cohen’s *d* effect sizes for subtype 1 vs. subtype 2 were found in SA (*d*=0.52 to 1.36) and SV (*d*=0.43 to 1.01) compared to CT (*d*=0.02 to 0.69). Cool colors = greater reductions in CT, SA, and SV in subtype 2 vs. subtype 1.

Resampling-based stability testing with 1000 iterations, each using a random sampling of 80% of the participants, revealed that the clusters were highly stable. On average, 92% of the time any given participant clustered with each other participant within their data-driven group, while only 8% of the time an individual was clustered with a participant in another group (Figure 2A, participant agreement matrix). The Adjusted Rand Index was 0.85 across 1000 iterations, indicating a high level of agreement between the clustering solutions considering chance agreement.

### Relative contribution of brain features in defining neuroanatomical subtypes of psychosis

In Figure 2B, cortical surface areas and subcortical volumes had higher NMI scores than thickness indicating that these features were relatively more important in determining the clusters (Table S6 for all feature NMI values). The top 10 model features, based on NMI scores, included subcortical volumes (hippocampus, amygdala, thalamus) and surface area of the medial orbitofrontal, inferior temporal, and precuneus. No thickness features appeared in the top 10. This remained the case when the NMI range was extended to the top 35 (Table S7). Resampling-based stability testing indicated that the top-ranking features based on NMI scores were consistent across iterations. On average, the top 10 variables remained among the top 35 more than 99% of the time across permutations (Figure S2).

ESs for the difference between subtypes were calculated for all brain features to further illustrate the relative importance of regional brain features that differentiate the subtypes. As shown in Figure 2C, and consistent with the NMI scores, ESs were very large (ES ∼1-1.5) for surface area, especially frontal, temporal, and midline cortical areas, and subcortical volumes, whereas ESs were considerably smaller for thickness.

### Neurostructural profiles of data-driven subtypes

Psychosis subtype neurostructural profiles were established by comparing each subtype to healthy individuals. In Figure 3, subtype 1 demonstrated small to medium (ES=0.16 to 0.52) enlargement of surface areas across most of the cortex, lower thickness in a single region of the prefrontal cortex, and larger basal ganglia volumes (caudate, putamen, globus pallidum). In contrast, subgroup 2 was characterized by widespread lower surface area and thickness, and smaller subcortical volumes. Effect sizes were notably larger for surface areas and subcortical volumes (ES=-0.33 to -1.00) relative to thickness (ES=-0.23 to -0.69). Complete results of the statistical analyses, including unthresholded statistical maps, are presented in the supplemental material.

**Figure 3.**
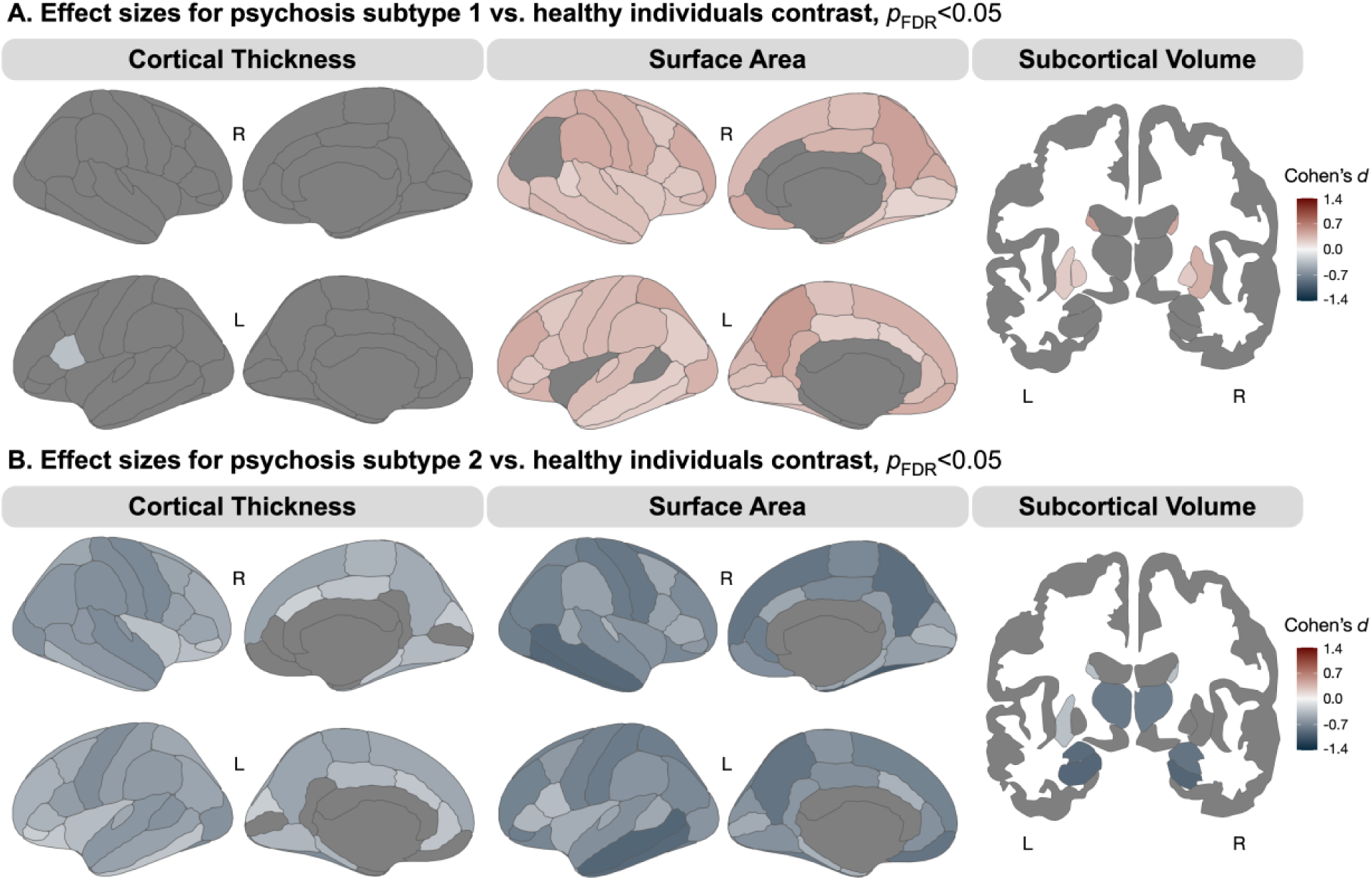
Data-driven psychosis subtypes demonstrate distinct neurostructural profiles compared to healthy individuals. Panel A: compared to healthy individuals, psychosis subtype 1 demonstrated small to moderate (*d*=0.16 to 0.52), but significantly larger surface area across the cortex and larger basal ganglia volumes. Panel B: compared to healthy individuals, psychosis subtype 2 demonstrated widespread lower surface area and cortical thickness, and smaller subcortical volumes. Effect sizes were larger for surface area and subcortical volume (*d*=-0.33 to -1.00) versus thickness (*d*=-0.23 to -0.69).

### Clinical characteristics of neuroanatomically-derived psychosis subtypes

Clinical characteristics and demographics of the two subtypes are presented in Table S5. The proportions of psychotic disorder illness stage were similar across the two subtypes, while proportions of SSD and BD diagnosis were different across the two subtypes (χ^2^(1, *N*=381)=9.93, *p*=0.002) (Figure 4A). There was no difference between subtypes in duration of illness and average dosage of antipsychotic medication (in CPZ equivalents).

**Figure 4.**
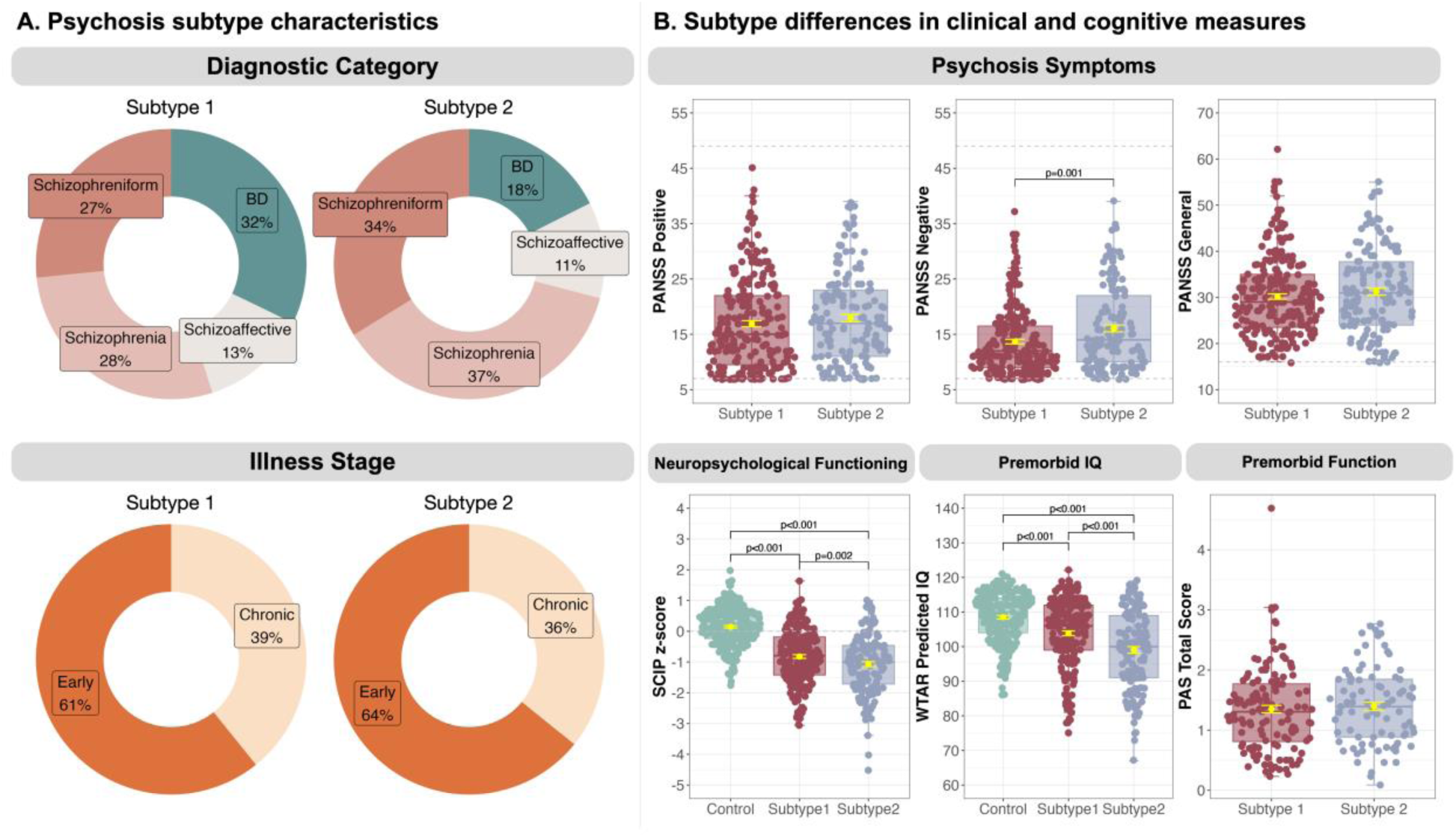
Clinical characteristics of neuroanatomically-derived psychosis subtypes. Panel A: psychosis subtypes displayed similar proportions of psychotic disorder diagnoses and illness stages. Panel B: subtype 2 displayed higher PANSS negative symptom scores compared to subtype 1, more severe cognitive impairment and lower premorbid intellectual functioning compared to both subtype 1 and healthy individuals.

Significant differences were observed between subtypes in severity of clinical symptoms and cognitive impairment (see Figure 4B, supplemental text). Subtype 2 exhibited higher PANSS negative symptom scores compared to subtype 1 (*b*=2.3, SE=0.7, *p*=0.001), and more severe cognitive impairment and lower premorbid intellectual functioning compared to both psychosis subtype 1 (*b*=-0.26, SE=0.08, *p*=0.002 and *b*=-4.91, SE=1.02, *p*<0.001, respectively) and healthy individuals (*b*=-1.22, SE=0.08, *p*<0.001 and *b*=-9.46, SE=1.01, *p*<0.001). PAS scores did not differ between subtypes.

### Multivariate cellular profiles of psychosis subtypes

We examined interregional associations between subtype neurostructural profiles (subtype vs. control effect size contrast maps), separately for surface area and thickness, with multiple cell-type abundances using PermCCA (Figure 5). Cell-type composite scores were significantly correlated with both surface area and thickness abnormalities in each subtype. Subtype 1 cortical thickness (*r*=0.79, *p*_FDR_=0.002) positively correlated with layer 4 intratelencephalic-projecting (L4 IT) excitatory neuron and negatively correlated with PAX6-expressing inhibitory interneuron abundance, meaning that cortical territories with more severe thinning overlap regions of greater PAX6 abundance but lower L4 IT. Subtype 2 cortical thickness (*r*=0.87, *p*_FDR_=0.002) positively correlated with oligodendrocyte (Oligo), SNCG-like inhibitory interneuron (SNCG), astrocyte (Astro), and somatostatin-expressing inhibitory interneuron (SST) and negatively correlated with parvalbumin-expressing inhibitory interneuron (PVALB) and vascular leptomeningeal cell (VLMC) abundance, meaning thinning is present in regions where PVALB and VLMC are abundant, but Oligo, Astro, SST, and SNCG are less prevalent. Subtype 1 surface area (*r*=0.69, *p*_FDR_=0.04) is positively correlated with PVALB and negatively correlated with layer 6 IT excitatory neuron (L6 IT Car3), Astro, and Oligo abundance, meaning regions of increased area overlap with those where PVALB is abundant, but L6 IT Car3, Astro, and Oligo are less prevalent. Subtype 2 surface area (*r*=0.72, *p*_FDR_=0.04) is negatively correlated with layer 5/6 near-projecting (L5/6 NP) and layer 5 IT (L5 IT) excitatory neuron abundance, meaning regions of thinner cortex overlap with those where L5/6 NP and L5 IT are most abundant.

**Figure 5.**
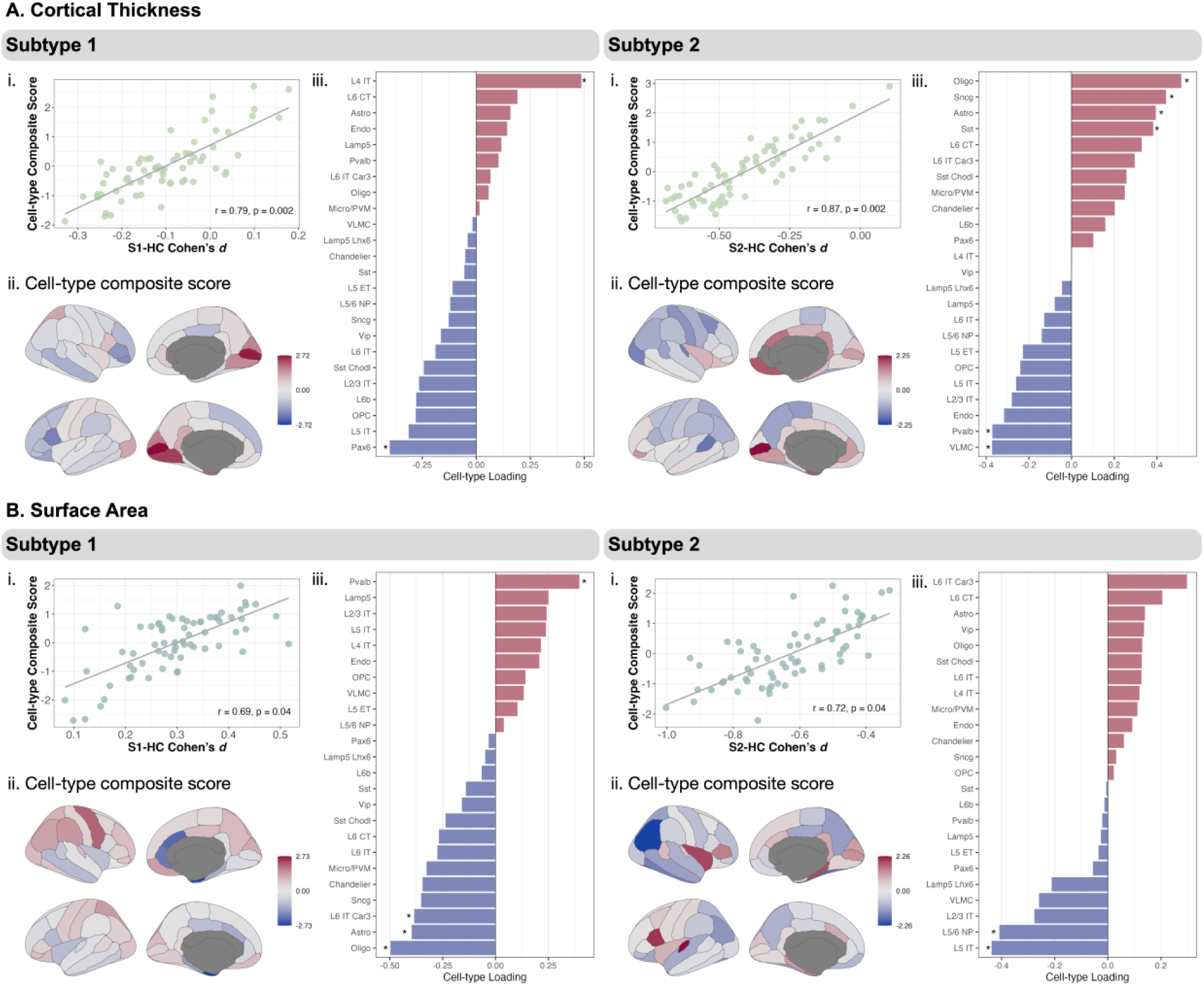
Multivariate cellular profiles follow spatial topography of neurostructural abnormalities in psychosis subtypes. Psychosis subtypes displayed differential spatial relationships between multiple cell-types and cortical thickness (panel A) and surface area (panel B) effect size maps (Cohen’s *d,* see Fig. 4). Panels A-B i: scatter plots for each subtype displaying the results of PermCCA, where the effect size map was positively related with a composite score of cell-type abundances (95% CI). Panels A-B ii: cell-type composite scores associated with each effect size map projected to the cortical surface. Panel A-B iii: canonical loadings of each cell-type to the composite score, significant cell-types indicated with an *, red = positive associations and blue = negative associations.

## DISCUSSION

We used a broad cohort of individuals with psychosis spanning early and chronic stages of both SSD and BD to assess the presence of neuroanatomical subtypes using a relatively new data-driven approach and integrated clinical and cell-type data to provide critical insight into potential diverging etiological pathways. Our findings show two distinct neuroanatomically-derived subtypes of psychosis driven by heterogeneity in cortical surface area, notably medial orbitofrontal and precuneus, and bilateral temporal and frontal regions, as well as subcortical volume of hippocampus, amygdala, and thalamus. Subtype 1 exhibited a small, but widespread increase in surface area and larger basal ganglia volumes compared to healthy individuals, whereas subtype 2 exhibited medium to large reductions in surface area and subcortical volumes. These findings align with a recent ENIGMA meta-and mega-analysis suggesting the greatest multimodal structural brain variability in schizophrenia for frontotemporal and subcortical structures (e.g., hippocampus, putamen) [47]. In contrast to the divergent pattern of abnormalities observed for surface area and subcortical volumes, lower cortical thickness was observed across much of the cortex in both subtypes but was considerably more severe and widespread in subtype 2. Subtypes were highly stable and not a consequence of demographic factors robustly associated with brain structure (e.g., age and sex) or illness chronicity; the proportion of early stage to chronic individuals was virtually identical in the two subtypes. The proportion of individuals with BD was relatively higher in subtype 1 (32% vs. 18% in subtype 2); however, approximately one quarter of BD individuals were in subtype 2 and the relative proportions of SSD diagnoses were similar across subtypes. Overall, the current findings provide strong support for two neuroanatomical subtypes that transcend illness stage and SSD and BD diagnoses.

Our findings are similar in several respects to prior data-driven approaches to parsing neurostructural heterogeneity in psychosis. Here, subtype 1 is consistent with findings by Chand et al. [21] showing a schizophrenia subtype marked by enlarged basal ganglia volumes [8, 21, 22]. A selective increase in striatal volume may be attributed to an excess of dopamine and dopamine receptors in schizophrenia [48, 49]. It has been speculated that for a subgroup of patients, this observed increase in D2 receptor binding may be genetically determined [50, 51] and persist in those even without antipsychotic treatment [8]. We further characterize this subtype by showing that larger basal ganglia volumes are accompanied by a modest, but extensive increase in surface area. Prior work focused exclusively on gross changes in cortical volume, meaning they were unable to distinguish between thickness and surface area. It is notable that expanded surface area in subtype 1 was observed in the context of cortical thinning that, while significant after correction for only one brain region, was present throughout the cortex. Given that cortical volume reflects the product of surface area and thickness, the combination of increased surface area and decreased thickness may explain why prior voxel-wise studies of grey matter volume did not detect this opposing pattern. Another dissociating aspect of our study is that our psychosis cohort included BD. Surface area abnormalities in SSD [7] and BD [13] are highly variable, with small effect sizes. This is especially the case for BD which has been linked to both regional increases and decreases [13]. It is noteworthy that 75% of BD patients included in the study clustered into subtype 1, however, this pattern of widespread thinner cortex and expanded surface area (vs. controls), notably in the precuneus and supramarginal gyrus, has been shown in non-deficit subgroups of schizophrenia, but not deficit subgroups [52]. Thus, subtype 1 may potentially capture a subset of patients more aligned with neurostructural patterns previously identified in BD and non-deficit subgroups of schizophrenia. Subtype 2 (39% of sample) in our psychosis cohort is consistent with findings showing widespread reductions in cortical volume and some subcortical regions [8, 21, 22]. We show that these reductions are observed in both surface area and thickness, and present in both SSD and BD. While a smaller portion of our sample, subtype 2 reflects the more typical case-control findings in SSD, but with much larger ESs (>0.50 vs. 0.20-0.40) suggesting a more impaired subtype of psychosis marked by prominent reductions in surface area, as well as thickness and subcortical volume. Consistent with greater neuroanatomical burden, subtype 2 displayed worse negative symptoms, and greater impairments in neuropsychological functioning and premorbid IQ compared to subtype 1.

Overlapping deficits in cortical thickness in the context of distinct abnormalities in surface area may indicate both shared and different mechanisms leading to structural abnormalities in subtypes 1 and 2. The synaptic pruning hypothesis posits that the normal developmental process of cortical thinning and remodeling that occurs from adolescence to young adulthood, when psychosis is most likely to emerge, is compromised in SSD [53]. This hypothesis is supported by longitudinal studies showing that the rate of cortical volume loss during this transitional period is higher in individuals at high-risk for psychosis that later transition to a psychotic disorder compared to both high-risk individuals that do not convert and healthy people [54]. Potentially the cause of this abnormality, whether it’s overactive synaptic pruning or a different mechanism(s), is shared across subtypes, but is more severe in subtype 2. In contrast, the striking differences in surface area strongly implies disparate trajectories in early brain development. Surface area increases dramatically from the 3^rd^ trimester until about 24-months postnatally after which it continues to increase, but at a much lower rate until peaking around age 11 [55, 56]. The development of surface area is complex and incompletely understood, involving neurogenesis and neuronal migration, including formation of cortical columns and relative proportion of basal radial glial cells which promote tangential spread of migrating neurons and cortical folding during fetal gestation [57–59]. Our findings suggest one or more of these processes is disrupted in a manner that promotes overgrowth in subtype 1 and undergrowth in subtype 2. Interestingly, both 22q11.2 deletion syndrome, which increases risk for psychosis approximately 30-fold, and copy number variants linked to developmental disorders are associated with smaller surface area, suggesting differences in genetic risk factors between subtypes [60, 61].

As cortical surface area and thickness display differential genetic architectures, we leveraged transcriptional data from the Allen Human Brain Atlas to examine how cell-type abundances spatially relate to subtype structural abnormalities. In both subtypes, greater abnormalities (expanded surface area in subtype 1, and greater cortical thinning in subtype 2) compared to healthy individuals were topographically linked with entorhinal/parahippocampus, anterior cingulate, and insula regions that display less glial cell (Oligo and Astro) abundance, and transverse temporal, parietal, and middle frontal regions that display greater PVALB abundance. While the present study cannot infer specific mechanisms, this finding emphasizes the potential functional relevance of glial cells and PVALB neurons in the divergent patterns of subtype structural abnormalities. A study examining transcriptome-MRI associations found marked heterogeneity in cell-type specific associations with cortical thickness in schizophrenia and suggests the presence of three cellular subtypes [11]. Here, subtype 2, aligns with a cellular subtype marked by severe and widespread cortical thinning present in brain regions where Astro is less prevalent and inhibitory (e.g., PVALB) neurons are more abundant. Psychosis subtype 1, potentially overlaps with a cellular subtype marked by milder cortical thinning, however, given the discrepancy in granularity of our cell-type analysis and Di Biase et al. [11] (24 vs. 7 cell-types), we are unable to discern whether the positive association we found with L4 IT and negative association with Pax6 aligns. Moving beyond the extant literature on thickness, we suggest that glial cells and PVALB, in a subset of individuals with psychosis, may differentially be related to expansions in surface area, again pointing to early diverging neurodevelopmental trajectories.

The largest effects of structural abnormalities across morphology measures and subtypes were found in surface area among subtype 2, compared to healthy controls. Smaller surface area in subtype 2 was topographically linked with parietal, superior temporal, and medial/lateral orbitofrontal regions that display greater L5 IT and L5/6 NP abundance. This finding emphasizes the potential relationship between layer 5 pyramidal neurons and surface area in the most impaired psychosis subtype. Layer 5 pyramidal neurons are critical for cortical input and output, where feedforward sensory information is integrated at their basal dendritic compartment, and feedback contextual information is integrated at their apical dendrites [62]. L5 IT neurons, located in superficial layer 5 (L5a), project intracortically and to the striatum, and receive primary input from higher-order thalamic nuclei [63]. These corticostriatal loops involving the prefrontal cortex, striatum, and thalamus, have long been hypothesized to be affected in schizophrenia and related to cognitive dysfunction [64, 65]. Spatial distributions of layer 5 IT neurons have also shown to map onto functional brain networks engaged in executive functions such as goal-oriented cognition (frontoparietal network), and visuospatial attention (dorsal attention), while co-expression of L5 IT and L5/6 IT neurons map onto a network engaged in recollection and mentalizing (default mode) processes [43]. As such, layer 5 pyramidal neurons, and their connectivity patterns may be related to the most severe abnormalities in brain structure and function among individuals with psychosis.

Strengths of the current work are the inclusion of a relatively large cohort of individuals with psychosis spanning early and chronic stages of both SSD and BD, use of a data-driven technique to parsing heterogeneity based on multiple brain morphology measures, rich phenotypic data to thoroughly characterize psychosis subtypes, and use of transcriptomics to develop cellular fingerprints of abnormalities in psychosis subtypes. Considering these strengths, several limitations of this study should be acknowledged. The data used was cross-sectional, and future longitudinal studies should examine if and how these psychosis subtypes change over time, and how that relates to clinical phenotypes. Second, while we conducted robust resampling-based stability testing of our clusters, the identified subtypes should be replicated in new samples to test the generalizability of the current study results. Third, the human microarray gene expression data obtained from bulk samples of six postmortem brains from the AHBA dataset were not from individuals living with SSD or BD. Future studies should examine how the identified subtype neurostructural abnormalities directly map onto cell-type distributions in people with psychotic disorders. Fourth, cell-type abundances are imputed from bulk-tissue microarray data that does not provide a direct estimate of gene transcription, but rather within-probe differences across samples. Bulk samples require aggregation into parcels to obtain robust estimates across donors, which limits the spatial resolution. Finally, an important future direction will be to examine multimodal biotypes, integrating functional and other structural (e.g., DTI) measures.

## Acknowledgments

This work was conducted in part using the resources of the Advanced Computing Center for Research and Education at Vanderbilt University, Nashville, TN.

## Funding

This work was supported by NIMH grants R01 MH102266, K24 MH126280, and P50 MH132642-5980 (awarded to NDW); R01 MH070560 (awarded to SH); R01 MH120080 (awarded to AJH), the Charlotte and Donald Test Fund and the Vanderbilt Institute for Clinical and Translational Research (through grant 1-UL-1-TR000445 from the National Center for Research Resources/NIH).

## Author Contributions

L.H.: conceptualization, methodology, analysis, visualization, original draft writing. X.Z.: transcriptomics conceptualization, methodology, analysis, visualization, and associated methods draft writing. B.R.: all imaging data storage and processing. A.S.H.: conceptualization, scientific and technical input, and imaging quality assurance. V.F., B.F.: collection, curation, and quality assurance of imaging and clinical data. S.H.: expert scientific and technical input, and funding acquisition. A.J.H.: transcriptomics conceptualization, methodology, expert scientific and technical input, and funding acquisition. N.W.: supervision, conceptualization, methodology, original draft writing, expert scientific and technical mentorship, and funding acquisition. All authors reviewed and revised the paper.

## Disclosures

No commercial support was received for the preparation of this manuscript, and the authors have no conflicts of interest to report.

## Notes

### Competing Interest Statement

The authors have declared no competing interest.

## REFERENCES

1. Calabrese, J. and Y. Al Khalili, Psychosis, in StatPearls. 2024: Treasure Island (FL).

2. De Peri, L., et al., Brain Structural Abnormalities at the Onset of Schizophrenia and Bipolar Disorder: A Meta-analysis of Controlled Magnetic Resonance Imaging Studies. Current Pharmaceutical Design, 2012. 18(4): p. 486–494.

3. Baker, J.T., et al., Disruption of cortical association networks in schizophrenia and psychotic bipolar disorder. JAMA Psychiatry, 2014. 71(2): p. 109–18.

4. Johnstone, E.C., et al., Cerebral ventricular size and cognitive impairment in chronic schizophrenia. Lancet, 1976. 2(7992): p. 924–6.

5. Shenton, M.E., T.J. Whitford, and M. Kubicki, Structural neuroimaging in schizophrenia: from methods to insights to treatments. Dialogues Clin Neurosci, 2010. 12(3): p. 317–32.

6. van Erp, T.G., et al., Subcortical brain volume abnormalities in 2028 individuals with schizophrenia and 2540 healthy controls via the ENIGMA consortium. Mol Psychiatry, 2016. 21(4): p. 585.

7. van Erp, T.G.M., et al., Cortical Brain Abnormalities in 4474 Individuals With Schizophrenia and 5098 Control Subjects via the Enhancing Neuro Imaging Genetics Through Meta Analysis (ENIGMA) Consortium. Biol Psychiatry, 2018. 84(9): p. 644–654.

8. Jiang, Y., et al., Neurostructural subgroup in 4291 individuals with schizophrenia identified using the subtype and stage inference algorithm. Nat Commun, 2024. 15(1): p. 5996.

9. Segal, A., et al., Regional, circuit and network heterogeneity of brain abnormalities in psychiatric disorders. Nat Neurosci, 2023. 26(9): p. 1613–1629.

10. Lv, J., et al., Individual deviations from normative models of brain structure in a large cross-sectional schizophrenia cohort. Mol Psychiatry, 2021. 26(7): p. 3512–3523.

11. Di Biase, M.A., et al., Cell type-specific manifestations of cortical thickness heterogeneity in schizophrenia. Molecular Psychiatry, 2022. 27(4): p. 2052–2060.

12. Madre, M., et al., Structural abnormality in schizophrenia versus bipolar disorder: A whole brain cortical thickness, surface area, volume and gyrification analyses. Neuroimage Clin, 2020. 25: p. 102131.

13. Matsumoto, J., et al., Cerebral cortical structural alteration patterns across four major psychiatric disorders in 5549 individuals. Mol Psychiatry, 2023. 28(11): p. 4915–4923.

14. Rashid, B. and V. Calhoun, Towards a brain-based predictome of mental illness. Hum Brain Mapp, 2020. 41(12): p. 3468–3535.

15. Chan, R.C., et al., Brain anatomical abnormalities in high-risk individuals, first-episode, and chronic schizophrenia: an activation likelihood estimation meta-analysis of illness progression. Schizophr Bull, 2011. 37(1): p. 177–88.

16. Adriano, F., C. Caltagirone, and G. Spalletta, Hippocampal volume reduction in first-episode and chronic schizophrenia: a review and meta-analysis. Neuroscientist, 2012. 18(2): p. 180–200.

17. Adriano, F., et al., Updated meta-analyses reveal thalamus volume reduction in patients with first-episode and chronic schizophrenia. Schizophr Res, 2010. 123(1): p. 1–14.

18. Clementz, B.A., et al., Identification of Distinct Psychosis Biotypes Using Brain-Based Biomarkers. Am J Psychiatry, 2016. 173(4): p. 373–84.

19. Clementz, B.A., et al., Psychosis Biotypes: Replication and Validation from the B-SNIP Consortium. Schizophr Bull, 2022. 48(1): p. 56–68.

20. Ivleva, E.I., et al., Brain Structure Biomarkers in the Psychosis Biotypes: Findings From the Bipolar-Schizophrenia Network for Intermediate Phenotypes. Biol Psychiatry, 2017. 82(1): p. 26–39.

21. Chand, G.B., et al., Two distinct neuroanatomical subtypes of schizophrenia revealed using machine learning. Brain, 2020. 143(3): p. 1027–1038.

22. Chand, G.B., et al., Schizophrenia Imaging Signatures and Their Associations With Cognition, Psychopathology, and Genetics in the General Population. American Journal of Psychiatry, 2022. 179(9): p. 650–660.

23. Panizzon, M.S., et al., Distinct genetic influences on cortical surface area and cortical thickness. Cereb Cortex, 2009. 19(11): p. 2728–35.

24. Wierenga, L.M., et al., Unique developmental trajectories of cortical thickness and surface area. Neuroimage, 2014. 87: p. 120–6.

25. Fortea, A., et al., Longitudinal Changes in Cortical Surface Area Associated With Transition to Psychosis in Adolescents at Clinical High Risk for the Disease. J Am Acad Child Adolesc Psychiatry, 2023. 62(5): p. 593–600.

26. Grasby, K.L., et al., The genetic architecture of the human cerebral cortex. Science, 2020. 367(6484).

27. Wang, B., et al., Similarity network fusion for aggregating data types on a genomic scale. Nat Methods, 2014. 11(3): p. 333–7.

28. Kay, S.R., A. Fiszbein, and L.A. Opler, The positive and negative syndrome scale (PANSS) for schizophrenia. Schizophr Bull, 1987. 13(2): p. 261–76.

29. Purdon, S., The screen for cognitive impairment in psychiatry (SCIP): administration manual and normative data. Edmonton, Alberta: PNL Inc., 2005.

30. Wechsler, D., Wechsler Test of Adult Reading: WTAR. Psychological Corporation, 2001.

31. Cannon-Spoor, H.E., S.G. Potkin, and R.J. Wyatt, Measurement of premorbid adjustment in chronic schizophrenia. Schizophr Bull, 1982. 8(3): p. 470–84.

32. Huo, Y., et al., Towards Portable Large-Scale Image Processing with High-Performance Computing. J Digit Imaging, 2018. 31(3): p. 304–314.

33. Harrigan, R.L., et al., Vanderbilt University Institute of Imaging Science Center for Computational Imaging XNAT: A multimodal data archive and processing environment. Neuroimage, 2016. 124(Pt B): p. 1097–1101.

34. Atkinson, D., et al., Automatic correction of motion artifacts in magnetic resonance images using an entropy focus criterion. Ieee Transactions on Medical Imaging, 1997. 16(6): p. 903–910.

35. Esteban, O., et al., MRIQC: Advancing the automatic prediction of image quality in MRI from unseen sites. Plos One, 2017. 12(9).

36. Fischl, B., FreeSurfer. Neuroimage, 2012. 62(2): p. 774–81.

37. Desikan, R.S., et al., An automated labeling system for subdividing the human cerebral cortex on MRI scans into gyral based regions of interest. Neuroimage, 2006. 31(3): p. 968–80.

38. Jacobs, G.R., et al., Integration of brain and behavior measures for identification of data-driven groups cutting across children with ASD, ADHD, or OCD. Neuropsychopharmacology, 2021. 46(3): p. 643–653.

39. Rand, W.M., Objective Criteria for Evaluation of Clustering Methods. Journal of the American Statistical Association, 1971. 66(336): p. 846–850.

40. Hubert, L. and P. Arabie, Comparing Partitions. Journal of Classification, 1985. 2(2-3): p. 193–218.

41. Benjamini, Y. and Y. Hochberg, Controlling the False Discovery Rate - a Practical and Powerful Approach to Multiple Testing. Journal of the Royal Statistical Society Series B-Statistical Methodology, 1995. 57(1): p. 289–300.

42. Nakagawa, S. and I.C. Cuthill, Effect size, confidence interval and statistical significance: a practical guide for biologists. Biological Reviews, 2007. 82(4): p. 591–605.

43. Zhang, X.H., et al., The cell-type underpinnings of the human functional cortical connectome. Nat Neurosci, 2025. 28(1): p. 150–160.

44. Jorstad, N.L., et al., Transcriptomic cytoarchitecture reveals principles of human neocortex organization. Science, 2023. 382(6667): p. eadf6812.

45. Hawrylycz, M.J., et al., An anatomically comprehensive atlas of the adult human brain transcriptome. Nature, 2012. 489(7416): p. 391–399.

46. Baum, G.L., et al., Development of structure-function coupling in human brain networks during youth. Proc Natl Acad Sci U S A, 2020. 117(1): p. 771–778.

47. Omlor, W., et al., Estimating Multimodal Structural Brain Variability in Schizophrenia Spectrum Disorders: A Worldwide ENIGMA. American Journal of Psychiatry, 2025. 182(4): p. 373–388.

48. Simpson, E.H., C. Kellendonk, and E. Kandel, A possible role for the striatum in the pathogenesis of the cognitive symptoms of schizophrenia. Neuron, 2010. 65(5): p. 585–96.

49. Kellendonk, C., et al., Transient and selective overexpression of dopamine D2 receptors in the striatum causes persistent abnormalities in prefrontal cortex functioning. Neuron, 2006. 49(4): p. 603–15.

50. Hirvonen, J., et al., Increased caudate dopamine D2 receptor availability as a genetic marker for schizophrenia. Archives of General Psychiatry, 2005. 62(4): p. 371–378.

51. Zvara, A., et al., Over-expression of dopamine D2 receptor and inwardly rectifying potassium channel genes in drug-naive schizophrenic peripheral blood lymphocytes as potential diagnostic markers. Dis Markers, 2005. 21(2): p. 61–9.

52. Banaj, N., et al., Cortical morphology in patients with the deficit and non-deficit syndrome of schizophrenia: a worldwide meta- and mega-analyses. Mol Psychiatry, 2023. 28(10): p. 4363–4373.

53. Feinberg, I., Schizophrenia: caused by a fault in programmed synaptic elimination during adolescence? J Psychiatr Res, 1982. 17(4): p. 319–34.

54. Cannon, T.D., et al., Progressive Reduction in Cortical Thickness as Psychosis Develops: A Multisite Longitudinal Neuroimaging Study of Youth at Elevated Clinical Risk. Biological Psychiatry, 2015. 77(2): p. 147–157.

55. Bethlehem, R.A.I., et al., Publisher Correction: Brain charts for the human lifespan. Nature, 2022. 610(7931): p. E6.

56. Huang, Y., et al., Mapping developmental regionalization and patterns of cortical surface area from 29 post-menstrual weeks to 2 years of age. Proc Natl Acad Sci U S A, 2022. 119(33): p. e2121748119.

57. Van Essen, D.C., Biomechanical models and mechanisms of cellular morphogenesis and cerebral cortical expansion and folding. Semin Cell Dev Biol, 2023. 140: p. 90–104.

58. Akula, S.K., D. Exposito-Alonso, and C.A. Walsh, Shaping the brain: The emergence of cortical structure and folding. Dev Cell, 2023. 58(24): p. 2836–2849.

59. Nowakowski, T.J., et al., Transformation of the Radial Glia Scaffold Demarcates Two Stages of Human Cerebral Cortex Development. Neuron, 2016. 91(6): p. 1219–1227.

60. Sun, D., et al., Large-scale mapping of cortical alterations in 22q11.2 deletion syndrome: Convergence with idiopathic psychosis and effects of deletion size. Mol Psychiatry, 2020. 25(8): p. 1822–1834.

61. Silva, A.I., et al., Penetrance of neurodevelopmental copy number variants is associated with variations in cortical morphology. Biological Psychiatry: Cognitive Neuroscience and Neuroimaging, 2025.

62. Takahashi, N., et al., Active dendritic currents gate descending cortical outputs in perception. Nat Neurosci, 2020. 23(10): p. 1277–1285.

63. Audette, N.J., et al., POm Thalamocortical Input Drives Layer-Specific Microcircuits in Somatosensory Cortex. Cereb Cortex, 2018. 28(4): p. 1312–1328.

64. Robbins, T.W., The case of frontostriatal dysfunction in schizophrenia. Schizophr Bull, 1990. 16(3): p. 391–402.

65. Shepherd, G.M., Corticostriatal connectivity and its role in disease. Nat Rev Neurosci, 2013. 14(4): p. 278–91.

