## Supplemental Information for "Subtyping psychotic disorders using a data-driven approach reveals divergent cortical and cellular signatures"

### SUPPLEMENTAL CONTENT

SUPPLEMENTAL TEXT

- Participant information
- MRI data acquisition
- Clinical characteristics of psychosis subtypes

SUPPLEMENTAL FIGURES

**Figure S1.** Sample inclusion/exclusion flow chart.

**Figure S2.** Stability of the top 35 brain features contributing to data-driven psychosis subtypes.

**Figure S3.** Unthresholded effect size brain maps of psychosis subtype 1 and 2 vs. healthy controls.

SUPPLEMENTAL TABLES

**Table S1.** Sample demographics and clinical characteristics by diagnostic category.

**Table S2.** Cortical thickness comparisons among patients (SSD and BD) and healthy controls.

**Table S3.** Surface area comparisons among patients (SSD and BD) and healthy controls.

**Table S4.** Subcortical volume comparisons among patients and healthy controls.

**Table S5.** Sample demographics and clinical characteristics by data-driven psychosis subtype.

**Table S6.** Normalized mutual information (NMI) values for all in-model brain features.

**Table S7.** Psychosis subtype comparisons on top 35 ranking model features contributing to participant similarity as determined by NMI.

**Table S8a.** Cortical thickness comparisons among data-driven subtype 1 and healthy controls.

**Table S8b.** Cortical thickness comparisons among data-driven subtype 2 and healthy controls.

**Table S9a.** Surface area comparisons among data-driven subtype 1 and healthy controls.

**Table S9b.** Surface area comparisons among data-driven subtype 2 and healthy controls.

**Table S10a.** Subcortical volume comparisons among data-driven subtype 1 and healthy controls.

**Table S10b.** Subcortical volume comparisons among data-driven subtype 2 and healthy controls.

REFERENCES

SUPPLEMENTAL TEXT

**Participant information.** Psychiatric diagnoses were confirmed, and ruled out in the case of healthy individuals, via interview with the Structured Clinical Interview for DSM-IV/V (SCID-IV/V) [1] conducted by trained research staff. See Supplemental Figure S1 for inclusion and exclusion criteria.

**MRI data acquisition.** Neuroimaging data were collected on two identical 3T Phillips Intera Achieva MRI scanners equipped with a 32-channel head coil located at Vanderbilt Institute for Imaging Sciences (VUIIS). In all three studies (CT00762866; R01MH070560; R01MH102266) a high resolution T1-weighted anatomical scan was collected for each individual with a 3D T1 fast field echo sequence with a 1mm$^3$ isotropic voxels (TR/TE = 8.0/3.7, FOV = 256 x 256 x 170, flip angle = 5°).

**Clinical characteristics of psychosis subtypes.** Subtype 2 exhibited significantly higher PANSS negative symptom scores compared to subtype 1 ($b$=2.3, SE=0.7, $p$=0.001), but no difference in positive or general symptomatology (positive: $b$=0.99, SE=0.87, $p$=0.25; general: $b$=1.05, SE=0.95, $p$=0.27). Higher negative symptoms indicate greater severity of symptoms including social and emotional withdrawal, and blunted affect. Regarding cognition, subtype 2 displayed significantly lower WTAR predicted IQ scores compared to subtype 1 ($b$=-4.91, SE=1.02, $p$<0.001) and healthy controls ($b$=-9.46, SE=1.01, $p$<0.001). Subtype 1 had significantly lower WTAR predicted IQ scores compared to healthy controls ($b$=-4.55, SE=0.88, $p$<0.001). Lower WTAR scores indicate worse premorbid intellectual functioning. Similarly, subtype 2 displayed significantly lower SCIP composite $z$-scores compared to subtype 1 ($b$=-0.26, SE=0.08, $p$=0.002) and healthy controls ($b$=-1.22, SE=0.08, $p$<0.001). Subtype 1 had significantly lower SCIP composite $z$-scores compared to healthy controls ($b$=-0.96, SE=0.07, $p$<0.001). Lower SCIP scores indicate worse current neuropsychological functioning. Subtype 1 and subtype 2 did not display differences on premorbid adjustment ($b$=0.03, SE=0.09, $p$=0.75).

SUPPLEMENTAL FIGURES

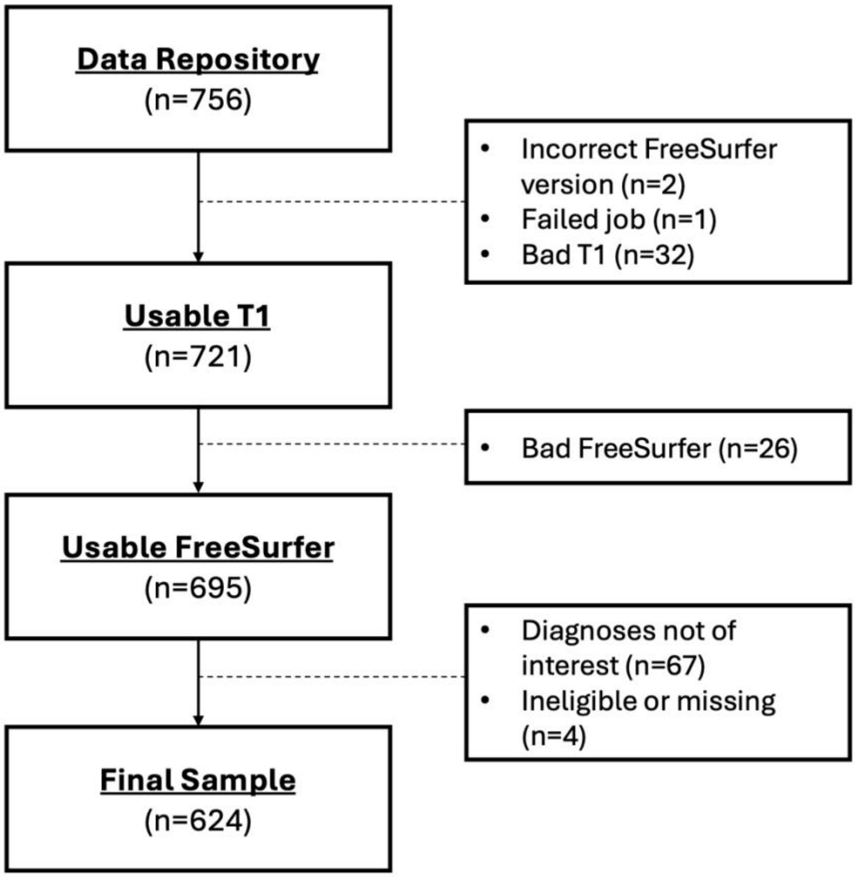


<u>**Figure S1.**</u> **Subject inclusion/exclusion flow chart**. A series of steps for the final selection of participants in the current study. The final sample of 624 individuals included 243 healthy controls, 101 people with bipolar disorder with psychotic features, and 280 people with schizophrenia spectrum disorder. See Table 1 and Table S1 for sample demographics.

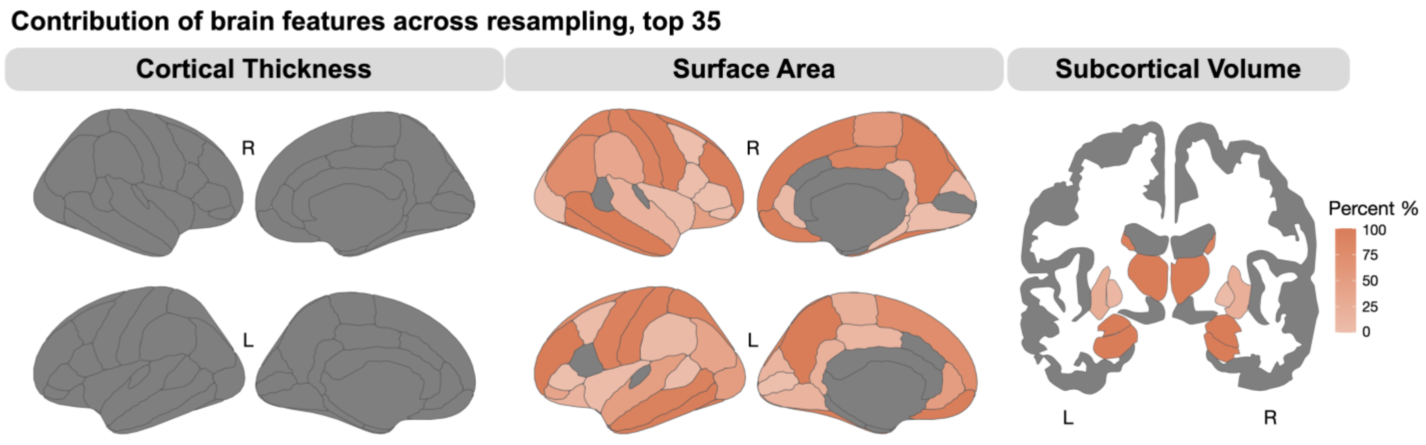


**Figure S2. Stability of the top 35 brain features contributing to data-driven psychosis subtypes**. The percentage of times that any given top contributing feature remained among the top 35 features across resampling was moderate to high.

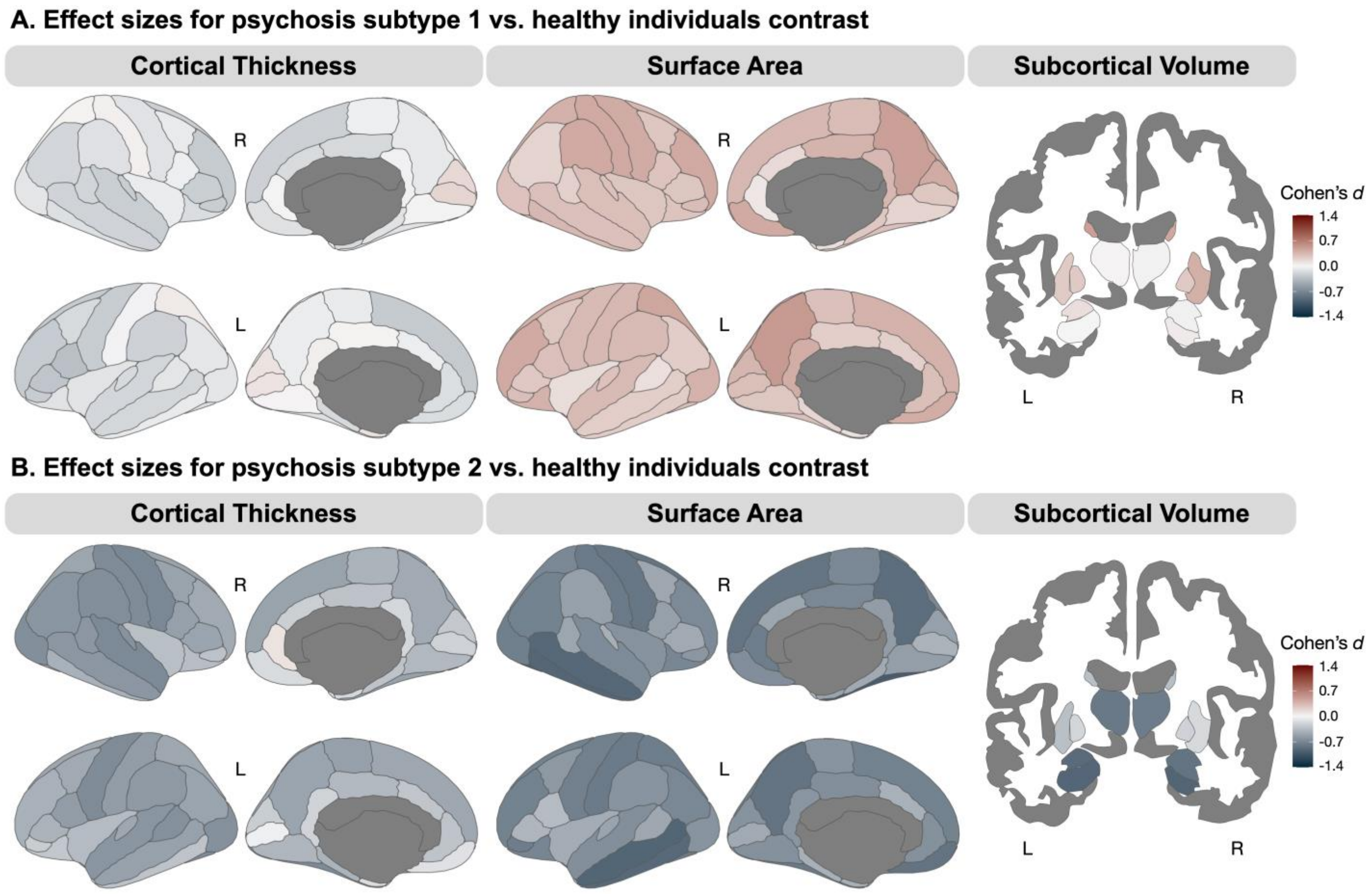


**<u>Figure S3.</u> Unthresholded effect size brain maps of psychosis subtype 1 and 2 vs. healthy controls.** Panel A: compared to healthy individuals, psychosis subtype 1 demonstrated patterns of primarily lower cortical thickness, larger surface area, and larger subcortical volumes. Panel B: compared to healthy individuals, psychosis subtype 2 demonstrated patterns of primarily lower cortical thickness and surface area, and smaller subcortical volumes.

SUPPLEMENTAL TABLES

**<u>Table S1.</u> Sample demographics and clinical characteristics by diagnostic category.**

| | **Healthy Individuals** (n=243) | **SSD** (n=280) | **BD** (n=101) | **Statistics** | | |
|---|---|---|---|---|---|---|
| | No. (%) | No. (%) | No. (%) | $\chi^2$ | df | *p*-value |
| **Sex** (female) | 91 (37.45) | 88 (31.43) | 49 (48.51) | 9.49 | 2 | *0.01* |
| **Race** | | | | 23.03 | 8 | *0.005* |
| White | 169 (69.55) | 172 (61.43) | 80 (79.21) | | | |
| Black | 54 (22.22) | 92 (32.86) | 12 (11.88) | | | |
| Asian | 11 (4.53) | 5 (1.78) | 3 (2.97) | | | |
| American Indian or Alaskan Native | 3 (1.23) | 2 (0.71) | 2 (1.98) | | | |
| Other | 6 (2.47) | 9 (3.22) | 4 (3.96) | | | |
| **Ethnicity** (Hispanic) | 12 (4.94) | 10 (3.57) | 7 (6.93) | 8.37 | 4 | 0.07 |
| **Handedness** (right) | 223 (91.77) | 249 (88.93) | 93 (92.08) | 1.25 | 2 | 0.53 |
| **Illness-Stage** (early/chronic) | --- | 179/101 | 57/43 | | | |
| | M ± SD | M ± SD | M ± SD | *F/t* | df | *p*-value |
| **Age** (years) | 28.03 ± 9.70 | 27.65 ± 10.32 | 30.36 ± 11.72 | 2.63 | 2 | 0.07 |
| **Education** (years) | 15.36 ± 2.11 | 13.41 ± 2.25 | 13.97 ± 1.94 | 52.28 | 2 | *<0.001 (C>BD/SSD)* |
| **Parental Education** (years) | 14.64 ± 2.29 | 14.63 ± 2.90 | 14.89 ± 2.26 | 0.37 | 2 | 0.69 |
| **Estimated Premorbid IQ** | 108.50 ± 7.57 | 100.88 ± 11.30 | 105.03 ± 9.05 | 39.28 | 2 | *<0.001 (C>BD>SSD)* |
| **SCIP** (z-score) | 0.14 ± 0.63 | -1 ± 0.94 | -0.67 ± 0.88 | 128.82 | 2 | *<0.001 (C>BD>SSD)* |
| **Age of Illness Onset** (years) | -- | 23.69 ± 8.45 | 21.00 ± 5.26 | 3.69 | 378 | *<0.001* |
| **Duration of Illness** (months) | --- | 74.71 ± 120.04 | 76.31 ± 108.68 | 0.12 | 378 | 0.91 |
| **PANSS Positive Symptoms** | --- | 18 ± 7.97 | 15.30 ± 8.93 | -2.77 | 372 | *0.005* |
| **PANSS Negative Symptoms** | --- | 15.96 ± 7.10 | 10.70 ± 3.87 | -6.91 | 372 | *<0.001* |
| **PANSS General Symptoms** | --- | 31.87 ± 9.21 | 26.98 ± 7.05 | -4.74 | 371 | *<0.001* |
| **Premorbid Adjustment** (total) | --- | 1.47 ± 0.67 | 1.09 ± 0.64 | -3.73 | 215 | *<0.001* |
| **CPZ Equivalents** | --- | 407.15 ± 551.70 | 292.01 ±210.26 | -1.78 | 318 | 0.08 |

**<u>Note.</u>** BD, bipolar with psychotic features; SSD, schizophrenia spectrum disorder; SCIP, Screen for Cognitive Impairment in Psychiatry; PANSS, Positive and Negative Syndrome Scale; Italics indicate significant *p*-values. Estimated premorbid IQ measured by the WTAR, Wechsler Test of Adult Reading predicted score. Duration of illness was defined as the time at which an individual first met criteria for psychosis (based on extensive interview, review of medical records, and collateral reports) until the date of study enrollment. CPZ, chlorpromazine equivalent.

**<u>Table S2.</u> Cortical thickness comparisons among patients and healthy controls.**

| Region | Cohen's *d* (PT v. HC) | SE | 95% CI | Estimate | *t* value | *p*-value | FDR *p*-value |
|---|---|---|---|---|---|---|---|
| **L banks STS** | **-0.294** | **0.0831** | **[-0.46, -0.13]** | **-0.04** | **-3.56** | **0.0004*** ** | **0.001** ** |
| **R banks STS** | **-0.341** | **0.0832** | **[-0.5, -0.18]** | **-0.05** | **-4.12** | **0.00004*** ** | **0.0003*** ** |
| L caudal anterior cingulate | -0.14 | 0.0827 | [-0.3, 0.02] | -0.03 | -1.70 | 0.09 | 0.11 |
| **R caudal anterior cingulate** | **-0.181** | **0.0828** | **[-0.34, -0.02]** | **-0.03** | **-2.19** | **0.03*** | **0.04*** |
| **L caudal middle frontal** | **-0.273** | **0.083** | **[-0.44, -0.11]** | **-0.04** | **-3.31** | **0.001** ** | **0.003** ** |
| **R caudal middle frontal** | **-0.231** | **0.0829** | **[-0.39, -0.07]** | **-0.03** | **-2.79** | **0.005** ** | **0.01*** |
| L cuneus | -0.0339 | 0.0826 | [-0.2, 0.13] | 0.00 | -0.41 | 0.68 | 0.72 |
| R cuneus | -0.01 | 0.0826 | [-0.17, 0.15] | 0.00 | -0.12 | 0.9 | 0.9 |
| L entorhinal | 0.0129 | 0.0826 | [-0.15, 0.18] | 0.00 | 0.16 | 0.88 | 0.9 |
| R entorhinal | -0.164 | 0.0828 | [-0.33, 0] | -0.05 | -1.98 | 0.048* | 0.07 |
| **L fusiform** | **-0.323** | **0.0831** | **[-0.49, -0.16]** | **-0.04** | **-3.91** | **0.0001** ** | **0.0004*** ** |
| **R fusiform** | **-0.263** | **0.083** | **[-0.43, -0.1]** | **-0.04** | **-3.18** | **0.002** ** | **0.004** ** |
| **L inferior parietal** | **-0.241** | **0.0829** | **[-0.4, -0.08]** | **-0.03** | **-2.92** | **0.004** ** | **0.008** ** |
| **R inferior parietal** | **-0.342** | **0.0832** | **[-0.51, -0.18]** | **-0.04** | **-4.13** | **0.00004*** ** | **0.0003*** ** |
| L inferior temporal | -0.153 | 0.0827 | [-0.32, 0.01] | -0.02 | -1.85 | 0.064 | 0.09 |
| **R inferior temporal** | **-0.239** | **0.0829** | **[-0.4, -0.08]** | **-0.03** | **-2.89** | **0.004** ** | **0.009** ** |
| L isthmus cingulate | -0.0503 | 0.0826 | [-0.21, 0.11] | -0.01 | -0.61 | 0.54 | 0.6 |
| R isthmus cingulate | -0.0825 | 0.0827 | [-0.24, 0.08] | -0.01 | -1.00 | 0.32 | 0.37 |
| **L lateral occipital** | **-0.293** | **0.0831** | **[-0.46, -0.13]** | **-0.03** | **-3.55** | **0.0004*** ** | **0.001** ** |
| **R lateral occipital** | **-0.296** | **0.0831** | **[-0.46, -0.13]** | **-0.03** | **-3.58** | **0.0004*** ** | **0.001** ** |
| **L lateral orbitofrontal** | **-0.175** | **0.0828** | **[-0.34, -0.01]** | **-0.03** | **-2.12** | **0.03*** | **0.049*** |
| **R lateral orbitofrontal** | **-0.216** | **0.0829** | **[-0.38, -0.05]** | **-0.03** | **-2.61** | **0.009** ** | **0.02*** |
| L lingual | -0.145 | 0.0827 | [-0.31, 0.02] | -0.02 | -1.75 | 0.08 | 0.11 |
| R lingual | -0.149 | 0.0827 | [-0.31, 0.01] | -0.02 | -1.81 | 0.07 | 0.09 |
| L medial orbitofrontal | -0.134 | 0.0827 | [-0.3, 0.03] | -0.02 | -1.62 | 0.11 | 0.13 |
| R medial orbitofrontal | -0.131 | 0.0827 | [-0.29, 0.03] | -0.01 | -1.59 | 0.11 | 0.13 |
| **L middle temporal** | **-0.308** | **0.0831** | **[-0.47, -0.15]** | **-0.04** | **-3.73** | **0.0002*** ** | **0.0008*** ** |
| **R middle temporal** | **-0.346** | **0.0832** | **[-0.51, -0.18]** | **-0.05** | **-4.19** | **0.00003*** ** | **0.0003*** ** |
| **L parahippocampal** | **-0.251** | **0.0829** | **[-0.41, -0.09]** | **-0.06** | **-3.04** | **0.003** ** | **0.006** ** |
| **R parahippocampal** | **-0.184** | **0.0828** | **[-0.35, -0.02]** | **-0.04** | **-2.23** | **0.03*** | **0.04*** |
| **L paracentral** | **-0.238** | **0.0829** | **[-0.4, -0.08]** | **-0.03** | **-2.88** | **0.004** ** | **0.009** ** |
| **R paracentral** | **-0.178** | **0.0828** | **[-0.34, -0.02]** | **-0.03** | **-2.16** | **0.03*** | **0.047*** |
| **L pars opercularis** | **-0.415** | **0.0835** | **[-0.58, -0.25]** | **-0.06** | **-5.02** | **<0.000001*** ** | **0.00002*** ** |
| **R pars opercularis** | **-0.223** | **0.0829** | **[-0.39, -0.06]** | **-0.03** | **-2.69** | **0.007** ** | **0.01*** |
| **L pars orbitalis** | **-0.246** | **0.0829** | **[-0.41, -0.08]** | **-0.05** | **-2.97** | **0.003** ** | **0.007** ** |
| **R pars orbitalis** | **-0.267** | **0.083** | **[-0.43, -0.1]** | **-0.05** | **-3.23** | **0.001** ** | **0.003** ** |
| **L pars triangularis** | **-0.337** | **0.0832** | **[-0.5, -0.17]** | **-0.05** | **-4.07** | **0.00005*** ** | **0.0003*** ** |
| **R pars triangularis** | **-0.337** | **0.0832** | **[-0.5, -0.17]** | **-0.05** | **-4.08** | **0.00005*** ** | **0.0003*** ** |
| L pericalcarine | 0.0498 | 0.0826 | [-0.11, 0.21] | 0.01 | 0.60 | 0.55 | 0.6 |
| R pericalcarine | 0.029 | 0.0826 | [-0.13, 0.19] | 0.00 | 0.35 | 0.73 | 0.76 |
| **L postcentral** | **-0.218** | **0.0829** | **[-0.38, -0.06]** | **-0.02** | **-2.64** | **0.009** ** | **0.02*** |
| **R postcentral** | **-0.224** | **0.0829** | **[-0.39, -0.06]** | **-0.02** | **-2.71** | **0.007** ** | **0.01*** |
| L posterior cingulate | -0.134 | 0.0827 | [-0.3, 0.03] | -0.02 | -1.62 | 0.11 | 0.13 |
| **R posterior cingulate** | **-0.223** | **0.0829** | **[-0.39, -0.06]** | **-0.03** | **-2.69** | **0.007** ** | **0.01*** |
| **L precentral** | **-0.401** | **0.0834** | **[-0.56, -0.24]** | **-0.05** | **-4.85** | **<0.000001*** ** | **0.00002*** ** |
| **R precentral** | **-0.324** | **0.0831** | **[-0.49, -0.16]** | **-0.04** | **-3.92** | **0.0001*** ** | **0.0004*** ** |
| **L precuneus** | **-0.218** | **0.0829** | **[-0.38, -0.06]** | **-0.03** | **-2.63** | **0.009** ** | **0.02*** |
| **R precuneus** | **-0.223** | **0.0829** | **[-0.39, -0.06]** | **-0.03** | **-2.69** | **0.007** ** | **0.01*** |
| **L rostral anterior cingulate** | **-0.199** | **0.0828** | **[-0.36, -0.04]** | **-0.03** | **-2.41** | **0.02*** | **0.03*** |
| R rostral anterior cingulate | 0.0445 | 0.0826 | [-0.12, 0.21] | 0.01 | 0.54 | 0.59 | 0.64 |
| **L rostral middle frontal** | **-0.318** | **0.0831** | **[-0.48, -0.16]** | **-0.04** | **-3.84** | **0.0001*** ** | **0.0005*** ** |
| **R rostral middle frontal** | **-0.334** | **0.0832** | **[-0.5, -0.17]** | **-0.04** | **-4.05** | **0.00006*** ** | **0.0003*** ** |
| **L superior frontal** | **-0.342** | **0.0832** | **[-0.51, -0.18]** | **-0.05** | **-4.14** | **0.00004*** ** | **0.0003*** ** |
| **R superior frontal** | **-0.33** | **0.0832** | **[-0.49, -0.17]** | **-0.05** | **-3.99** | **0.00007*** ** | **0.0003*** ** |
| L superior parietal | -0.144 | 0.0827 | [-0.31, 0.02] | -0.02 | -1.74 | 0.08 | 0.11 |

| | | | | | | | |
|---|---|---|---|---|---|---|---|
| **R superior parietal** | **-0.178** | **0.0828** | **[-0.34, -0.02]** | **-0.02** | **-2.16** | **0.03*** | **0.047*** |
| **L superior temporal** | **-0.342** | **0.0832** | **[-0.51, -0.18]** | **-0.05** | **-4.14** | **0.00004***** | **0.0003***** |
| **R superior temporal** | **-0.402** | **0.0834** | **[-0.57, -0.24]** | **-0.06** | **-4.86** | **<0.000001***** | **0.00002***** |
| **L supramarginal** | **-0.349** | **0.0832** | **[-0.51, -0.19]** | **-0.04** | **-4.22** | **0.00003***** | **0.0003***** |
| **R supramarginal** | **-0.315** | **0.0831** | **[-0.48, -0.15]** | **-0.04** | **-3.81** | **0.0002***** | **0.0006***** |
| **L frontal pole** | **-0.275** | **0.083** | **[-0.44, -0.11]** | **-0.07** | **-3.33** | **0.0009***** | **0.003**** |
| R frontal pole | -0.0107 | 0.0826 | [-0.17, 0.15] | 0.00 | -0.13 | 0.89 | 0.9 |
| L temporal pole | -0.0777 | 0.0827 | [-0.24, 0.08] | -0.02 | -0.94 | 0.35 | 0.39 |
| R temporal pole | -0.0798 | 0.0827 | [-0.24, 0.08] | -0.03 | -0.97 | 0.33 | 0.39 |
| **L transverse temporal** | **-0.283** | **0.083** | **[-0.45, -0.12]** | **-0.06** | **-3.43** | **0.0007***** | **0.002**** |
| **R transverse temporal** | **-0.383** | **0.0834** | **[-0.55, -0.22]** | **-0.08** | **-4.63** | **<0.000001***** | **0.00002***** |
| **L insula** | **-0.197** | **0.0828** | **[-0.36, -0.03]** | **-0.03** | **-2.39** | **0.02*** | **0.03*** |
| R insula | -0.159 | 0.0828 | [-0.32, 0] | -0.03 | -1.93 | 0.05 | 0.08 |

**Note.** General linear models per region controlling for age, sex, scan version, and T1w quality (MRIQC efc). *$p<0.05$, **$p<0.01$, ***$p<0.001$. FDR corrected using Benjamini-Hochberg procedure across all CT regions. Regions with $p_{FDR}<0.05$ are depicted in bold. PT = patient, HC = healthy control, SE = standard error, L = left, R = right.

**Table S3.** **Surface area comparisons among patients and healthy controls.**

| Region | Cohen's *d* (PT v. HC) | SE | 95% CI | Estimate | *t* value | *p*-value | FDR *p*-value |
|---|---|---|---|---|---|---|---|
| L banks STS | -0.107 | 0.0827 | [-0.27, 0.06] | -17.71 | -1.29 | 0.197 | 0.974 |
| R banks STS | -0.0596 | 0.0827 | [-0.22, 0.1] | -7.71 | -0.72 | 0.471 | 0.979 |
| L caudal anterior cingulate | -0.0353 | 0.0826 | [-0.2, 0.13] | -4.71 | -0.43 | 0.669 | 0.979 |
| R caudal anterior cingulate | -0.0917 | 0.0827 | [-0.25, 0.07] | -13.82 | -1.11 | 0.268 | 0.974 |
| L caudal middle frontal | -0.0626 | 0.0827 | [-0.23, 0.1] | -22.13 | -0.76 | 0.449 | 0.979 |
| R caudal middle frontal | -0.0183 | 0.0826 | [-0.18, 0.14] | -6.46 | -0.22 | 0.825 | 0.979 |
| L cuneus | -0.0138 | 0.0826 | [-0.18, 0.15] | -2.94 | -0.17 | 0.867 | 0.979 |
| R cuneus | 0.051 | 0.0826 | [-0.11, 0.21] | 10.72 | 0.62 | 0.537 | 0.979 |
| L entorhinal | -0.102 | 0.0827 | [-0.26, 0.06] | -9.31 | -1.23 | 0.217 | 0.974 |
| R entorhinal | -0.116 | 0.0827 | [-0.28, 0.05] | -9.39 | -1.40 | 0.161 | 0.974 |
| L fusiform | -0.114 | 0.0827 | [-0.28, 0.05] | -40.92 | -1.38 | 0.168 | 0.974 |
| R fusiform | -0.184 | 0.0828 | [-0.35, -0.02] | -60.18 | -2.23 | 0.026* | 0.442 |
| L inferior parietal | -0.12 | 0.0827 | [-0.28, 0.04] | -71.37 | -1.45 | 0.148 | 0.974 |
| R inferior parietal | -0.158 | 0.0828 | [-0.32, 0.004] | -111.73 | -1.91 | 0.056 | 0.762 |
| L inferior temporal | -0.189 | 0.0828 | [-0.35, -0.03] | -89.15 | -2.28 | 0.023* | 0.442 |
| R inferior temporal | -0.129 | 0.0827 | [-0.29, 0.03] | -54.62 | -1.56 | 0.119 | 0.974 |
| L isthmus cingulate | 0.0188 | 0.0826 | [-0.14, 0.18] | 2.88 | 0.23 | 0.820 | 0.979 |
| R isthmus cingulate | 0.0048 | 0.0826 | [-0.16, 0.17] | 0.63 | 0.06 | 0.954 | 0.986 |
| L lateral occipital | -0.0028 | 0.0826 | [-0.17, 0.16] | -1.65 | -0.03 | 0.973 | 0.986 |
| R lateral occipital | -0.106 | 0.0827 | [-0.27, 0.06] | -64.97 | -1.28 | 0.200 | 0.974 |
| L lateral orbitofrontal | -0.0411 | 0.0826 | [-0.2, 0.12] | -12.59 | -0.50 | 0.619 | 0.979 |
| R lateral orbitofrontal | -0.0947 | 0.0827 | [-0.26, 0.07] | -32.29 | -1.15 | 0.252 | 0.974 |
| L lingual | -0.0768 | 0.0827 | [-0.24, 0.09] | -30.56 | -0.93 | 0.353 | 0.979 |
| R lingual | -0.0878 | 0.0827 | [-0.25, 0.07] | -36.26 | -1.06 | 0.288 | 0.979 |
| L medial orbitofrontal | -0.0652 | 0.0827 | [-0.23, 0.1] | -14.34 | -0.79 | 0.430 | 0.979 |
| R medial orbitofrontal | 0.0128 | 0.0826 | [-0.15, 0.18] | 2.78 | 0.16 | 0.877 | 0.979 |
| L middle temporal | -0.209 | 0.0828 | [-0.37, -0.05] | -85.59 | -2.53 | 0.012* | 0.394 |
| R middle temporal | -0.151 | 0.0827 | [-0.31, 0.01] | -66.62 | -1.83 | 0.067 | 0.763 |
| L parahippocampal | -0.0559 | 0.0826 | [-0.22, 0.11] | -4.00 | -0.68 | 0.499 | 0.979 |
| R parahippocampal | -0.0271 | 0.0826 | [-0.19, 0.14] | -2.07 | -0.33 | 0.743 | 0.979 |
| L paracentral | -0.0495 | 0.0826 | [-0.21, 0.11] | -8.04 | -0.60 | 0.549 | 0.979 |
| R paracentral | -0.0626 | 0.0827 | [-0.23, 0.1] | -11.20 | -0.76 | 0.449 | 0.979 |
| L pars opercularis | 0.0466 | 0.0826 | [-0.12, 0.21] | 11.79 | 0.56 | 0.573 | 0.979 |
| R pars opercularis | -0.0358 | 0.0826 | [-0.2, 0.13] | -7.15 | -0.43 | 0.665 | 0.979 |
| L pars orbitalis | -0.0965 | 0.0827 | [-0.26, 0.07] | -9.11 | -1.17 | 0.243 | 0.974 |
| R pars orbitalis | -0.0312 | 0.0826 | [-0.19, 0.13] | -3.49 | -0.38 | 0.706 | 0.979 |
| L pars triangularis | 0.0015 | 0.0826 | [-0.16, 0.16] | 0.28 | 0.02 | 0.986 | 0.986 |
| R pars triangularis | -0.0127 | 0.0826 | [-0.18, 0.15] | -2.88 | -0.15 | 0.878 | 0.979 |
| L pericalcarine | 0.0141 | 0.0826 | [-0.15, 0.18] | 3.60 | 0.17 | 0.865 | 0.979 |
| R pericalcarine | 0.0027 | 0.0826 | [-0.16, 0.17] | 0.70 | 0.03 | 0.974 | 0.986 |
| L postcentral | -0.0834 | 0.0827 | [-0.25, 0.08] | -37.99 | -1.01 | 0.313 | 0.979 |
| R postcentral | -0.0318 | 0.0826 | [-0.19, 0.13] | -13.91 | -0.38 | 0.701 | 0.979 |
| L posterior cingulate | -0.111 | 0.0827 | [-0.27, 0.05] | -18.15 | -1.34 | 0.181 | 0.974 |
| R posterior cingulate | -0.0341 | 0.0826 | [-0.2, 0.13] | -5.72 | -0.41 | 0.680 | 0.979 |
| L precentral | -0.0762 | 0.0827 | [-0.24, 0.09] | -39.21 | -0.92 | 0.357 | 0.979 |
| R precentral | -0.0476 | 0.0826 | [-0.21, 0.12] | -22.51 | -0.58 | 0.564 | 0.979 |
| L precuneus | -0.0052 | 0.0826 | [-0.17, 0.16] | -2.42 | -0.06 | 0.950 | 0.986 |
| R precuneus | -0.0296 | 0.0826 | [-0.19, 0.13] | -14.41 | -0.36 | 0.720 | 0.979 |
| L rostral anterior cingulate | -0.0501 | 0.0826 | [-0.21, 0.11] | -8.23 | -0.61 | 0.545 | 0.979 |
| R rostral anterior cingulate | -0.23 | 0.0829 | [-0.39, -0.07] | -29.46 | -2.78 | 0.006** | 0.374 |
| L rostral middle frontal | -0.0178 | 0.0826 | [-0.18, 0.14] | -12.31 | -0.22 | 0.829 | 0.979 |
| R rostral middle frontal | -0.002 | 0.0826 | [-0.16, 0.16] | -1.49 | -0.02 | 0.981 | 0.986 |
| L superior frontal | -0.0454 | 0.0826 | [-0.21, 0.12] | -37.45 | -0.55 | 0.583 | 0.979 |
| R superior frontal | -0.0908 | 0.0827 | [-0.25, 0.07] | -74.66 | -1.10 | 0.272 | 0.974 |
| L superior parietal | -0.0144 | 0.0826 | [-0.18, 0.15] | -9.90 | -0.17 | 0.861 | 0.979 |

| | | | | | | | |
|---|---|---|---|---|---|---|---|
| R superior parietal | -0.0289 | 0.0826 | [-0.19, 0.13] | -17.99 | -0.35 | 0.727 | 0.979 |
| L superior temporal | -0.0425 | 0.0826 | [-0.21, 0.12] | -18.97 | -0.51 | 0.607 | 0.979 |
| R superior temporal | -0.0608 | 0.0827 | [-0.22, 0.1] | -23.60 | -0.74 | 0.462 | 0.979 |
| L supramarginal | -0.0356 | 0.0826 | [-0.2, 0.13] | -22.82 | -0.43 | 0.667 | 0.979 |
| R supramarginal | 0.0551 | 0.0826 | [-0.11, 0.22] | 27.20 | 0.67 | 0.505 | 0.979 |
| L frontal pole | 0.0303 | 0.0826 | [-0.13, 0.19] | 1.06 | 0.37 | 0.714 | 0.979 |
| R frontal pole | -0.0181 | 0.0826 | [-0.18, 0.14] | -0.81 | -0.22 | 0.827 | 0.979 |
| L temporal pole | 0.0021 | 0.0826 | [-0.16, 0.16] | 0.14 | 0.03 | 0.980 | 0.986 |
| R temporal pole | -0.0467 | 0.0826 | [-0.21, 0.12] | -3.24 | -0.56 | 0.572 | 0.979 |
| L transverse temporal | -0.0163 | 0.0826 | [-0.18, 0.15] | -1.06 | -0.20 | 0.844 | 0.979 |
| R transverse temporal | 0.0132 | 0.0826 | [-0.15, 0.18] | 0.60 | 0.16 | 0.873 | 0.979 |
| L insula | -0.0956 | 0.0827 | [-0.26, 0.07] | -26.64 | -1.16 | 0.248 | 0.974 |
| R insula | -0.0685 | 0.0827 | [-0.23, 0.09] | -15.78 | -0.83 | 0.407 | 0.979 |

**Note.** General linear models per region controlling for age, sex, scan version, and T1w quality (MRIQC efc). *$p<0.05$, **$p<0.01$, ***$p<0.001$. FDR corrected using Benjamini-Hochberg procedure across all SA regions. Regions with $p_{FDR}<0.05$ are depicted in bold. PT = patient, HC = healthy control, SE = standard error, L = left, R = right.

**<u>Table S4.</u> Subcortical volume comparisons among patients and healthy controls.**

| Region | Cohen's *d* (PT v. HC) | SE | 95% CI | Estimate | *t* value | *p*-value | FDR *p*-value |
|---|---|---|---|---|---|---|---|
| **L thalamus** | **-0.291** | **0.0831** | **[-0.45, -0.13]** | **-164.67** | **-3.52** | **0.00047*** ** | **0.0013** ** |
| **R thalamus** | **-0.298** | **0.0831** | **[-0.46, 0.14]** | **-154.61** | **-3.60** | **0.00034*** ** | **0.0012** ** |
| L caudate | 0.168 | 0.0828 | [0.01, 0.33] | 67.01 | 2.03 | 0.04 | 0.062 |
| R caudate | 0.139 | 0.0827 | [-0.02, 0.30] | 56.32 | 1.68 | 0.09 | 0.13 |
| L putamen | 0.021 | 0.0826 | [-0.41, 0.18] | 12.82 | 0.26 | 0.79 | 0.79 |
| R putamen | 0.18 | 0.0828 | [0.02, 0.34] | 80.82 | 2.18 | 0.03* | 0.06 |
| L pallidum | 0.08 | 0.0827 | [-0.08, 0.24] | 16.97 | 0.97 | 0.33 | 0.39 |
| R pallidum | 0.071 | 0.0827 | [-0.09, 0.23] | 13.39 | 0.86 | 0.39 | 0.42 |
| **L hippocampus** | **-0.339** | **0.0832** | **[-0.50, -0.18]** | **-111.44** | **-4.10** | **0.00005*** ** | **0.0007*** ** |
| **R hippocampus** | **-0.303** | **0.0831** | **[-0.47, -0.14]** | **-100.27** | **-3.67** | **0.00027*** ** | **0.0012** ** |
| **L amygdala** | **-0.246** | **0.0829** | **[-0.41, -0.08]** | **-50.87** | **-2.97** | **0.003** ** | **0.007** ** |
| **R amygdala** | **-0.322** | **0.0831** | **[-0.49, -0.16]** | **-68.49** | **-3.89** | **0.0001*** ** | **0.0007*** ** |
| L accumbens | -0.169 | 0.0828 | [-0.33, -0.01] | -17.35 | -2.04 | 0.04* | 0.062 |
| R accumbens | -0.116 | 0.0827 | [-0.28, 0.05] | -10.24 | -1.39 | 0.16 | 0.20 |

**<u>Note.</u>** General linear models per region controlling for age, sex, scan version, T1w quality (MRIQC efc), and intracranial volume. **p*<0.05, ***p*<0.01, ****p*<0.001. FDR corrected using Benjamini-Hochberg procedure across all SV regions. Regions with $p_{FDR}$<0.05 are depicted in bold. PT = patient, HC = healthy control, SE = standard error, L = left, R = right.

**<u>Table S5.</u> Sample demographics and clinical characteristics by data-driven psychosis subtype.**

| | **Healthy Individuals** (n=243) | **Subtype 1** (n=233) | **Subtype 2** (n=148) | **Statistics** | | |
|---|---|---|---|---|---|---|
| | No. (%) | No. (%) | No. (%) | $\chi^2$ | df | *p*-value |
| **Sex** (female) | 91 (37.45) | 87 (37.34) | 50 (33.78) | 0.64 | 2 | 0.73 |
| **Race** | | | | 57.65 | 8 | *<0.001* |
| White | 169 (69.55) | 185 (79.40) | 67 (45.27) | | | |
| Black | 54 (22.22) | 36 (15.45) | 68 (45.95) | | | |
| Asian | 11 (4.53) | 2 (0.89) | 6 (4.05) | | | |
| American Indian or Alaskan Native | 3 (1.23) | 3 (1.29) | 1 (0.68) | | | |
| Other | 6 (2.47) | 7 (3.00) | 6 (4.05) | | | |
| **Ethnicity** (Hispanic) | 12 (4.94) | 13 (5.58) | 4 (2.70) | 8.05 | 4 | 0.09 |
| **Handedness** (right) | 223 (91.77) | 214 (91.85) | 128 (86.49) | 4.09 | 2 | 0.13 |
| **Diagnosis** (SSD/BD) | --- | 158/75 | 122/26 | 9.93 | 1 | *0.002* |
| **Illness-Stage** (early/chronic) | --- | 141/91 | 95/53 | 0.31 | 1 | 0.58 |
| | M ± SD | M ± SD | M ± SD | *F/t* | df | *p*-value |
| **Age** (years) | 28.03 ± 9.70 | 28.66 ± 11.06 | 27.90 ± 10.30 | 0.32 | 2 | 0.72 |
| **Education** (years) | 15.36 ± 2.11 | 13.60 ± 2.20 | 13.47 ± 2.16 | 49.73 | 2 | *<0.001 (C>S1/S2)* |
| **Parental Education** (years) | 14.64 ± 2.29 | 14.83 ± 2.75 | 14.49 ± 2.76 | 0.71 | 2 | 0.49 |
| **Estimated Premorbid IQ** | 108.50 ± 7.57 | 103.86 ± 10.17 | 99.04 ± 11.37 | 44.87* | 2 | *<0.001* |
| **SCIP** (*z*-score) | 0.14 ± 0.63 | -0.82 ± 0.89 | -1.07 ± 0.98 | 137.23* | 2 | *<0.001* |
| **Age of Illness Onset** (years) | --- | 21.72 ± 6.61 | 21.70 ± 5.95 | 0.04 | 378 | 0.97 |
| **Duration of Illness** (months) | --- | 79.28 ± 121.55 | 68.62 ± 109.62 | 0.87 | 378 | 0.39 |
| **PANSS Positive Symptoms** | --- | 16.91 ± 8.32 | 17.93 ± 8.25 | 1.14* | 373 | 0.25 |
| **PANSS Negative Symptoms** | --- | 13.67 ± 6.17 | 16.05 ± 7.52 | 3.27* | 373 | *0.001* |
| **PANSS General Symptoms** | --- | 30.19 ± 8.75 | 31.28 ± 9.26 | 1.11* | 372 | 0.27 |
| **Premorbid Adjustment** (total) | --- | 1.35 ± 0.71 | 1.40 ± 0.64 | 0.32* | 216 | 0.75 |
| **CPZ Equivalents** | --- | 400.08 ± 588.37 | 343.67 ± 251.19 | 0.98 | 318 | 0.33 |

**<u>Note.</u>** SSD, Schizophrenia Spectrum Disorder; BD, Bipolar with Psychotic Features. Duration of illness was defined as the time at which an individual first met criteria for psychosis (based on extensive interview, review of medical records, and collateral reports) until the date of study enrollment. One patient in subtype 1 missing illness-stage data. Group differences on estimated premorbid IQ measured by the WTAR, Wechsler Test of Adult Reading predicted score; SCIP, Screen for Cognitive Impairment in Psychiatry; PANSS, Positive and Negative Syndrome Scale; and Premorbid Adjustment (n=217) were tested using separate general linear models (denoted with *, *t*- and *F*-values using the *aov* package in R [version 4.4.1] displayed here for consistency), detailed in the main text and Figure 4. Italics indicate significant *p*-values.

**Table S6. Normalized mutual information (NMI) values for all in-model brain features.**

| Rank | Model Brain Feature | NMI | Rank | Model Brain Feature | NMI |
|---|---|---|---|---|---|
| 1 | R hippocampus | 0.258 | 76 | L frontal pole, SA | 0.045 |
| 2 | L amygdala | 0.219 | 77 | L pars opercularis, SA | 0.043 |
| 3 | L hippocampus | 0.212 | 78 | L pars triangularis, SA | 0.039 |
| 4 | R amygdala | 0.194 | 79 | L entorhinal, SA | 0.039 |
| 5 | L medial orbitofrontal, SA | 0.191 | 80 | R pericalcarine, SA | 0.039 |
| 6 | R inferior temporal, SA | 0.188 | 81 | L temporal pole, SA | 0.038 |
| 7 | L precuneus, SA | 0.186 | 82 | R temporal pole, SA | 0.038 |
| 8 | R precuneus, SA | 0.185 | 83 | R precentral, CT | 0.036 |
| 9 | L precentral, SA | 0.177 | 84 | R pericalcarine, CT | 0.035 |
| 10 | L thalamus | 0.174 | 85 | L lingual, CT | 0.034 |
| 11 | R fusiform, SA | 0.169 | 86 | R frontal pole, SA | 0.034 |
| 12 | R thalamus | 0.166 | 87 | R inferior parietal, CT | 0.032 |
| 13 | R middle temporal, SA | 0.163 | 88 | R caudal middle frontal, CT | 0.032 |
| 14 | L inferior temporal, SA | 0.160 | 89 | R lateral occipital, CT | 0.031 |
| 15 | L superior parietal, SA | 0.155 | 90 | L caudal anterior cingulate, SA | 0.031 |
| 16 | L postcentral, SA | 0.152 | 91 | L postcentral, CT | 0.031 |
| 17 | R posterior cingulate, SA | 0.152 | 92 | L precuneus, CT | 0.029 |
| 18 | L caudate | 0.150 | 93 | L posterior cingulate, CT | 0.029 |
| 19 | L rostral middle frontal, SA | 0.149 | 94 | L banks sts, CT | 0.029 |
| 20 | R rostral middle frontal, SA | 0.149 | 95 | R cuneus, CT | 0.028 |
| 21 | L middle temporal, SA | 0.145 | 96 | R supramarginal, CT | 0.026 |
| 22 | L superior frontal, SA | 0.143 | 97 | L inferior parietal, CT | 0.026 |
| 23 | L superior temporal, SA | 0.142 | 98 | L fusiform, CT | 0.023 |
| 24 | R medial orbitofrontal, SA | 0.141 | 99 | R pars opercularis, CT | 0.022 |
| 25 | R superior parietal, SA | 0.140 | 100 | R lingual, CT | 0.021 |
| 26 | R caudate | 0.139 | 101 | L superior parietal, CT | 0.021 |
| 27 | L fusiform, SA | 0.139 | 102 | R superior parietal, CT | 0.019 |
| 28 | R superior frontal, SA | 0.135 | 103 | L cuneus, CT | 0.019 |
| 29 | R precentral, SA | 0.129 | 104 | R precuneus, CT | 0.019 |
| 30 | L putamen | 0.126 | 105 | R paracentral, CT | 0.019 |
| 31 | R postcentral, SA | 0.126 | 106 | R superior temporal, CT | 0.018 |
| 32 | R supramarginal, SA | 0.125 | 107 | L superior temporal, CT | 0.017 |
| 33 | L lateral orbitofrontal, SA | 0.122 | 108 | L paracentral, CT | 0.016 |
| 34 | L pars orbitalis, SA | 0.119 | 109 | L precentral, CT | 0.015 |
| 35 | R superior temporal, SA | 0.116 | 110 | R fusiform, CT | 0.015 |
| 36 | R paracentral, SA | 0.116 | 111 | R superior frontal, CT | 0.013 |
| 37 | R accumbens | 0.116 | 112 | L middle temporal, CT | 0.012 |
| 38 | R putamen | 0.115 | 113 | R insula, CT | 0.011 |
| 39 | L inferior parietal, SA | 0.113 | 114 | R inferior temporal, CT | 0.011 |
| 40 | L paracentral, SA | 0.111 | 115 | R transverse temporal, CT | 0.010 |
| 41 | R lateral orbitofrontal, SA | 0.110 | 116 | L rostral middle frontal, CT | 0.010 |
| 42 | R inferior parietal, SA | 0.110 | 117 | R middle temporal, CT | 0.009 |
| 43 | L lateral occipital, SA | 0.107 | 118 | L isthmus cingulate, CT | 0.009 |
| 44 | L posterior cingulate, SA | 0.106 | 119 | L transverse temporal, CT | 0.009 |
| 45 | L rostral anterior cingulate, SA | 0.105 | 120 | R rostral middle frontal, CT | 0.008 |
| 46 | L cuneus, SA | 0.102 | 121 | R banks sts, CT | 0.008 |
| 47 | L isthmus cingulate, SA | 0.098 | 122 | L caudal anterior cingulate, CT | 0.008 |
| 48 | R pars orbitalis, SA | 0.097 | 123 | R lateral orbitofrontal, CT | 0.007 |
| 49 | L transverse temporal, SA | 0.096 | 124 | R isthmus cingulate, CT | 0.006 |
| 50 | R isthmus cingulate, SA | 0.091 | 125 | R pars triangularis, CT | 0.006 |
| 51 | L lingual, SA | 0.091 | 126 | L caudal middle frontal, CT | 0.005 |
| 52 | L supramarginal, SA | 0.091 | 127 | R posterior cingulate, CT | 0.005 |
| 53 | L insula, SA | 0.089 | 128 | L superior frontal, CT | 0.005 |

| | | | | | |
|---|---|---|---|---|---|
| 54 | L pallidum | 0.086 | 129 | L inferior temporal, CT | 0.004 |
| 55 | R rostral anterior cingulate, SA | 0.083 | 130 | L supramarginal, CT | 0.003 |
| 56 | R lateral occipital, SA | 0.083 | 131 | L insula, CT | 0.003 |
| 57 | L caudal middle frontal, SA | 0.082 | 132 | R parahippocampal, CT | 0.003 |
| 58 | R insula, SA | 0.080 | 133 | L entorhinal, CT | 0.002 |
| 59 | R pars triangularis, SA | 0.079 | 134 | L pars orbitalis, CT | 0.002 |
| 60 | R banks sts, SA | 0.079 | 135 | L lateral orbitofrontal, CT | 0.002 |
| 61 | L pericalcarine, SA | 0.077 | 136 | L medial orbitofrontal, CT | 0.001 |
| 62 | R pars opercularis, SA | 0.076 | 137 | R medial orbitofrontal, CT | 0.001 |
| 63 | R pallidum | 0.075 | 138 | L pericalcarine, CT | 0.001 |
| 64 | R lingual, SA | 0.075 | 139 | R frontal pole, CT | 0.001 |
| 65 | R cuneus, SA | 0.071 | 140 | L pars opercularis, CT | 0.001 |
| 66 | R caudal middle frontal, SA | 0.070 | 141 | R temporal pole, CT | 0.001 |
| 67 | L accumbens | 0.069 | 142 | L rostral anterior cingulate, CT | 0.000 |
| 68 | R transverse temporal, SA | 0.067 | 143 | R caudal anterior cingulate, CT | 0.000 |
| 69 | L lateral occipital, CT | 0.057 | 144 | L parahippocampal, CT | 0.000 |
| 70 | L banks sts, SA | 0.054 | 145 | R pars orbitalis, CT | 0.000 |
| 71 | L parahippocampal, SA | 0.051 | 146 | L frontal pole, CT | 0.000 |
| 72 | R caudal anterior cingulate, SA | 0.051 | 147 | R rostral anterior cingulate, CT | 0.000 |
| 73 | R parahippocampal, SA | 0.050 | 148 | L pars triangularis, CT | 0.000 |
| 74 | R postcentral, CT | 0.050 | 149 | R entorhinal, CT | 0.000 |
| 75 | R entorhinal, SA | 0.045 | 150 | L temporal pole, CT | 0.000 |

**Note.** SA, surface area; CT, cortical thickness; sts, superior temporal sulcus.

**Table S7.** Psychosis subtype comparisons on top 35 ranking model features contributing to participant similarity as determined by NMI.

| No. | Top Contributing Features | NMI | Contrast | Cohen's *d* | SE | 95% CI | *t*-value | *p*FDR |
|---|---|---|---|---|---|---|---|---|
| 1 | R hippocampus | 0.258 | S1-HC | 0.057 | 0.09 | [-0.13,0.24] | 0.61 | 0.697 |
| | | | **S2-HC** | **-0.929** | **0.11** | **[-1.15,-0.71]** | **-8.47** | **2.07e-15** |
| 2 | L amygdala | 0.219 | S1-HC | 0.128 | 0.09 | [-0.06,0.31] | 1.36 | 0.242 |
| | | | **S2-HC** | **-0.887** | **0.11** | **[-1.11,-0.67]** | **-8.08** | **2.05e-14** |
| 3 | L hippocampus | 0.212 | S1-HC | -0.002 | 0.09 | [-0.19,0.18] | -0.03 | 0.979 |
| | | | **S2-HC** | **-0.926** | **0.11** | **[-1.15,-0.7]** | **-8.44** | **2.07e-15** |
| 4 | R amygdala | 0.194 | S1-HC | -0.027 | 0.09 | [-0.21,0.16] | -0.29 | 0.927 |
| | | | **S2-HC** | **-0.830** | **0.11** | **[-1.05,-0.61]** | **-7.57** | **6.35e-13** |
| 5 | L medial orbitofrontal, SA | 0.191 | **S1-HC** | **0.405** | **0.09** | **[0.22,0.59]** | **4.40** | **1.87e-05** |
| | | | **S2-HC** | **-0.829** | **0.11** | **[-1.04,-0.62]** | **-7.90** | **8.28e-14** |
| 6 | R inferior temporal, SA | 0.188 | **S1-HC** | **0.346** | **0.09** | **[0.16,0.53]** | **3.76** | **2.32e-04** |
| | | | **S2-HC** | **-0.919** | **0.11** | **[-1.13,-0.71]** | **-8.77** | **3.09e-16** |
| 7 | L precuneus, SA | 0.186 | **S1-HC** | **0.516** | **0.09** | **[0.33,0.7]** | **5.60** | **6.46e-08** |
| | | | **S2-HC** | **-0.833** | **0.11** | **[-1.04,-0.62]** | **-7.95** | **6.88e-14** |
| 8 | R precuneus, SA | 0.185 | **S1-HC** | **0.492** | **0.09** | **[0.31,0.68]** | **5.34** | **2.54e-07** |
| | | | **S2-HC** | **-0.866** | **0.11** | **[-1.08,-0.66]** | **-8.26** | **7.80e-15** |
| 9 | L precentral, SA | 0.177 | **S1-HC** | **0.332** | **0.09** | **[0.15,0.51]** | **3.60** | **3.95e-04** |
| | | | **S2-HC** | **-0.739** | **0.11** | **[-0.95,-0.53]** | **-7.05** | **1.61e-11** |
| 10 | L thalamus | 0.174 | S1-HC | 0.003 | 0.09 | [-0.18,0.19] | 0.03 | 0.979 |
| | | | **S2-HC** | **-0.793** | **0.11** | **[-1.01,-0.57]** | **-7.22** | **5.42e-12** |
| 11 | R fusiform, SA | 0.169 | **S1-HC** | **0.297** | **0.09** | **[0.12,0.48]** | **3.23** | **0.001** |
| | | | **S2-HC** | **-1.003** | **0.11** | **[-1.22,-0.79]** | **-9.57** | **1.56e-18** |
| 12 | R thalamus | 0.166 | S1-HC | -0.021 | 0.09 | [-0.21,0.16] | -0.22 | 0.927 |
| | | | **S2-HC** | **-0.770** | **0.11** | **[-0.99,-0.55]** | **-7.02** | **1.80e-11** |
| 13 | R middle temporal, SA | 0.163 | **S1-HC** | **0.3** | **0.09** | **[0.12,0.48]** | **3.26** | **0.001** |
| | | | **S2-HC** | **-0.907** | **0.11** | **[-1.12,-0.7]** | **-8.65** | **5.76e-16** |
| 14 | L inferior temporal, SA | 0.160 | **S1-HC** | **0.233** | **0.09** | **[0.05,0.42]** | **2.53** | **0.012** |
| | | | **S2-HC** | **-0.902** | **0.11** | **[-1.11,-0.69]** | **-8.60** | **6.84e-16** |
| 15 | L superior parietal, SA | 0.155 | **S1-HC** | **0.452** | **0.09** | **[0.27,0.64]** | **4.91** | **2.11e-06** |
| | | | **S2-HC** | **-0.758** | **0.11** | **[-0.97,-0.55]** | **-7.23** | **4.99e-12** |
| 16 | L postcentral, SA | 0.152 | **S1-HC** | **0.345** | **0.09** | **[0.16,0.53]** | **3.75** | **2.34e-04** |
| | | | **S2-HC** | **-0.783** | **0.11** | **[-0.99,-0.57]** | **-7.46** | **1.26e-12** |
| 17 | R posterior cingulate, SA | 0.152 | **S1-HC** | **0.373** | **0.09** | **[0.19,0.56]** | **4.04** | **7.68e-05** |
| | | | **S2-HC** | **-0.686** | **0.11** | **[-0.9,-0.48]** | **-6.54** | **3.22e-10** |
| 18 | L caudate | 0.150 | **S1-HC** | **0.486** | **0.1** | **[0.3,0.67]** | **5.17** | **8.02e-07** |
| | | | **S2-HC** | **-0.329** | **0.11** | **[-0.55,-0.11]** | **-3.00** | **0.005** |
| 19 | L rostral middle frontal, SA | 0.149 | **S1-HC** | **0.427** | **0.09** | **[0.24,0.61]** | **4.63** | **6.95e-06** |
| | | | **S2-HC** | **-0.727** | **0.11** | **[-0.94,-0.52]** | **-6.93** | **3.02e-11** |
| 20 | R rostral middle frontal, SA | 0.149 | **S1-HC** | **0.425** | **0.09** | **[0.24,0.61]** | **4.62** | **7.31e-06** |
| | | | **S2-HC** | **-0.68** | **0.11** | **[-0.89,-0.47]** | **-6.48** | **4.38e-10** |
| 21 | L middle temporal, SA | 0.145 | **S1-HC** | **0.215** | **0.09** | **[0.03,0.4]** | **2.33** | **0.02** |
| | | | **S2-HC** | **-0.93** | **0.11** | **[-1.14,-0.72]** | **-8.87** | **2.02e-16** |
| 22 | L superior frontal, SA | 0.143 | **S1-HC** | **0.381** | **0.09** | **[0.2,0.56]** | **4.14** | **5.48e-05** |
| | | | **S2-HC** | **-0.732** | **0.11** | **[-0.94,-0.52]** | **-6.98** | **2.37e-11** |
| 23 | L superior temporal, SA | 0.142 | **S1-HC** | **0.332** | **0.09** | **[0.15,0.51]** | **3.61** | **3.95e-04** |
| | | | **S2-HC** | **-0.644** | **0.11** | **[-0.85,-0.44]** | **-6.14** | **3.00e-09** |
| 24 | R medial orbitofrontal, SA | 0.141 | **S1-HC** | **0.433** | **0.09** | **[0.25,0.62]** | **4.70** | **5.29e-06** |
| | | | **S2-HC** | **-0.651** | **0.11** | **[-0.86,-0.44]** | **-6.21** | **2.10e-09** |
| 25 | R superior parietal, SA | 0.140 | **S1-HC** | **0.386** | **0.09** | **[0.2,0.57]** | **4.19** | **4.57e-05** |
| | | | **S2-HC** | **-0.692** | **0.11** | **[-0.9,-0.48]** | **-6.60** | **2.30e-10** |
| 26 | R caudate | 0.139 | **S1-HC** | **0.44** | **0.1** | **[0.25,0.63]** | **4.69** | **7.70e-06** |
| | | | **S2-HC** | **-0.336** | **0.11** | **[-0.55,-0.12]** | **-3.06** | **0.005** |
| 27 | L fusiform, SA | 0.139 | **S1-HC** | **0.295** | **0.09** | **[0.11,0.48]** | **3.20** | **0.001** |
| | | | **S2-HC** | **-0.787** | **0.11** | **[-1,-0.58]** | **-7.50** | **1.03e-12** |

| | | | | | | | | |
|---|---|---|---|---|---|---|---|---|
| 28 | R superior frontal, SA | 0.135 | **S1-HC** | **0.354** | **0.09** | **[0.17,0.54]** | **3.84** | **1.72e-04** |
| | | | **S2-HC** | **-0.818** | **0.11** | **[-1.03,-0.61]** | **-7.80** | **1.52e-13** |
| 29 | R precentral, SA | 0.129 | **S1-HC** | **0.423** | **0.09** | **[0.24,0.61]** | **4.60** | **7.77e-06** |
| | | | **S2-HC** | **-0.807** | **0.11** | **[-1.02,-0.6]** | **-7.70** | **2.84e-13** |
| 30 | L putamen | 0.126 | **S1-HC** | **0.24** | **0.09** | **[0.05,0.43]** | **2.55** | **0.016** |
| | | | **S2-HC** | **-0.33** | **0.11** | **[-0.55,-0.11]** | **-3.01** | **0.005** |
| 31 | R postcentral, SA | 0.126 | **S1-HC** | **0.433** | **0.09** | **[0.25,0.62]** | **4.70** | **5.29e-06** |
| | | | **S2-HC** | **-0.778** | **0.11** | **[-0.99,-0.57]** | **-7.42** | **1.61e-12** |
| 32 | R supramarginal, SA | 0.125 | **S1-HC** | **0.432** | **0.09** | **[0.25,0.61]** | **4.69** | **5.52e-06** |
| | | | **S2-HC** | **-0.533** | **0.11** | **[-0.74,-0.33]** | **-5.08** | **9.12e-07** |
| 33 | L lateral orbitofrontal, SA | 0.122 | **S1-HC** | **0.378** | **0.09** | **[0.2,0.56]** | **4.10** | **6.17e-05** |
| | | | **S2-HC** | **-0.714** | **0.11** | **[-0.92,-0.51]** | **-6.81** | **6.26e-11** |
| 34 | L pars orbitalis, SA | 0.119 | **S1-HC** | **0.31** | **0.09** | **[0.13,0.49]** | **3.37** | **8.82e-04** |
| | | | **S2-HC** | **-0.762** | **0.11** | **[-0.97,-0.55]** | **-7.27** | **4.20e-12** |
| 35 | R superior temporal, SA | 0.116 | **S1-HC** | **0.314** | **0.09** | **[0.13,0.5]** | **3.41** | **7.71e-04** |
| | | | **S2-HC** | **-0.666** | **0.11** | **[-0.88,-0.46]** | **-6.36** | **9.15e-10** |

**Note.** Data-driven psychosis subtype comparisons with healthy individuals using separate generalized linear models while including age, sex, scan version, and T1 image quality metric efc as covariates. Subcortical models additionally controlled for intracranial volume (ICV). FDR corrected using Benjamini-Hochberg procedure, regions with $p_{FDR}<0.05$ are depicted in bold. S1, subtype 1; S2, subtype 2; HC, healthy control; SA, surface area; R, right; L, left.

**Table S8a. Cortical thickness comparisons among data-driven subtype 1 and healthy controls.**

| Region | Cohen's *d* (S1 v. HC) | SE | 95% CI | Estimate | *t* value | *p*-value | FDR *p*-value |
|---|---|---|---|---|---|---|---|
| L banks STS | -0.090 | 0.092 | [-0.27,0.09] | -0.013 | -0.98 | 0.3272 | 0.5070 |
| R banks STS | -0.190 | 0.092 | [-0.37,-0.01] | -0.029 | -2.06 | 0.0400 | 0.1699 |
| L caudal anterior cingulate | -0.045 | 0.092 | [-0.23,0.14] | -0.008 | -0.49 | 0.6245 | 0.7864 |
| R caudal anterior cingulate | -0.150 | 0.092 | [-0.33,0.03] | -0.025 | -1.62 | 0.1048 | 0.2639 |
| L caudal middle frontal | -0.157 | 0.092 | [-0.34,0.02] | -0.021 | -1.70 | 0.0891 | 0.2396 |
| R caudal middle frontal | -0.058 | 0.092 | [-0.24,0.12] | -0.008 | -0.63 | 0.5285 | 0.7047 |
| L cuneus | 0.096 | 0.092 | [-0.09,0.28] | 0.013 | 1.04 | 0.2981 | 0.4826 |
| R cuneus | 0.156 | 0.092 | [-0.03,0.34] | 0.019 | 1.69 | 0.0916 | 0.2396 |
| L entorhinal | 0.100 | 0.092 | [-0.08,0.28] | 0.031 | 1.09 | 0.2764 | 0.4698 |
| R entorhinal | -0.157 | 0.092 | [-0.34,0.02] | -0.050 | -1.70 | 0.0895 | 0.2396 |
| L fusiform | -0.117 | 0.092 | [-0.3,0.06] | -0.013 | -1.27 | 0.2039 | 0.4078 |
| R fusiform | -0.107 | 0.092 | [-0.29,0.07] | -0.014 | -1.16 | 0.2475 | 0.4435 |
| L inferior parietal | -0.083 | 0.092 | [-0.26,0.1] | -0.010 | -0.90 | 0.3660 | 0.5410 |
| R inferior parietal | -0.187 | 0.092 | [-0.37,-0.01] | -0.022 | -2.03 | 0.0425 | 0.1699 |
| L inferior temporal | -0.063 | 0.092 | [-0.24,0.12] | -0.008 | -0.68 | 0.4946 | 0.6780 |
| R inferior temporal | -0.124 | 0.092 | [-0.31,0.06] | -0.017 | -1.35 | 0.1783 | 0.3910 |
| L isthmus cingulate | 0.041 | 0.092 | [-0.14,0.22] | 0.006 | 0.45 | 0.6537 | 0.8070 |
| R isthmus cingulate | -0.022 | 0.092 | [-0.2,0.16] | -0.003 | -0.24 | 0.8125 | 0.9057 |
| L lateral occipital | -0.089 | 0.092 | [-0.27,0.09] | -0.010 | -0.96 | 0.3355 | 0.5070 |
| R lateral occipital | -0.090 | 0.092 | [-0.27,0.09] | -0.010 | -0.97 | 0.3305 | 0.5070 |
| L lateral orbitofrontal | -0.111 | 0.092 | [-0.29,0.07] | -0.016 | -1.20 | 0.2297 | 0.4435 |
| R lateral orbitofrontal | -0.139 | 0.092 | [-0.32,0.04] | -0.017 | -1.51 | 0.1321 | 0.3097 |
| L lingual | 0.006 | 0.092 | [-0.18,0.19] | 0.001 | 0.07 | 0.9456 | 0.9598 |
| R lingual | -0.012 | 0.092 | [-0.19,0.17] | -0.001 | -0.13 | 0.8964 | 0.9378 |
| L medial orbitofrontal | -0.143 | 0.092 | [-0.32,0.04] | -0.018 | -1.55 | 0.1225 | 0.2975 |
| R medial orbitofrontal | -0.122 | 0.092 | [-0.3,0.06] | -0.014 | -1.32 | 0.1865 | 0.3963 |
| L middle temporal | -0.180 | 0.092 | [-0.36,0.002] | -0.025 | -1.95 | 0.0518 | 0.1826 |
| R middle temporal | -0.201 | 0.092 | [-0.38,-0.02] | -0.027 | -2.18 | 0.0294 | 0.1335 |
| L parahippocampal | -0.157 | 0.092 | [-0.34,0.02] | -0.039 | -1.70 | 0.0891 | 0.2396 |
| R parahippocampal | -0.109 | 0.092 | [-0.29,0.07] | -0.022 | -1.18 | 0.2373 | 0.4435 |
| L paracentral | -0.062 | 0.092 | [-0.24,0.12] | -0.009 | -0.68 | 0.4985 | 0.6780 |
| R paracentral | -0.036 | 0.092 | [-0.22,0.15] | -0.005 | -0.39 | 0.6960 | 0.8070 |
| **L pars opercularis** | **-0.329** | **0.093** | **[-0.51,-0.15]** | **-0.049** | **-3.58** | **0.0004** | **0.0256** |
| R pars opercularis | -0.036 | 0.092 | [-0.22,0.15] | -0.005 | -0.39 | 0.7002 | 0.8070 |
| L pars orbitalis | -0.247 | 0.092 | [-0.43,-0.07] | -0.046 | -2.68 | 0.0076 | 0.0688 |
| R pars orbitalis | -0.239 | 0.092 | [-0.42,-0.06] | -0.045 | -2.59 | 0.0097 | 0.0688 |
| L pars triangularis | -0.288 | 0.093 | [-0.47,-0.11] | -0.043 | -3.13 | 0.0019 | 0.0631 |
| R pars triangularis | -0.236 | 0.092 | [-0.42,-0.05] | -0.036 | -2.56 | 0.0107 | 0.0688 |
| L pericalcarine | 0.099 | 0.092 | [-0.08,0.28] | 0.014 | 1.07 | 0.2852 | 0.4730 |
| R pericalcarine | 0.178 | 0.092 | [-0.003,0.36] | 0.023 | 1.93 | 0.0537 | 0.1826 |
| L postcentral | -0.016 | 0.092 | [-0.2,0.17] | -0.002 | -0.17 | 0.8645 | 0.9378 |
| R postcentral | 0.036 | 0.092 | [-0.15,0.22] | 0.004 | 0.39 | 0.6986 | 0.8070 |
| L posterior cingulate | 0.013 | 0.092 | [-0.17,0.19] | 0.002 | 0.14 | 0.8895 | 0.9378 |
| R posterior cingulate | -0.168 | 0.092 | [-0.35,0.01] | -0.021 | -1.82 | 0.0689 | 0.2232 |
| L precentral | -0.248 | 0.092 | [-0.43,-0.07] | -0.032 | -2.69 | 0.0074 | 0.0688 |
| R precentral | -0.107 | 0.092 | [-0.29,0.07] | -0.013 | -1.16 | 0.2478 | 0.4435 |
| L precuneus | -0.046 | 0.092 | [-0.23,0.14] | -0.005 | -0.50 | 0.6170 | 0.7864 |
| R precuneus | -0.074 | 0.092 | [-0.26,0.11] | -0.009 | -0.81 | 0.4197 | 0.6073 |
| L rostral anterior cingulate | -0.135 | 0.092 | [-0.32,0.05] | -0.024 | -1.47 | 0.1423 | 0.3227 |
| R rostral anterior cingulate | 0.007 | 0.092 | [-0.17,0.19] | 0.001 | 0.08 | 0.9372 | 0.9598 |
| L rostral middle frontal | -0.255 | 0.092 | [-0.44,-0.07] | -0.030 | -2.77 | 0.0058 | 0.0688 |
| R rostral middle frontal | -0.254 | 0.092 | [-0.44,-0.07] | -0.031 | -2.75 | 0.0061 | 0.0688 |
| L superior frontal | -0.256 | 0.092 | [-0.44,-0.07] | -0.035 | -2.78 | 0.0056 | 0.0688 |
| R superior frontal | -0.221 | 0.092 | [-0.4,-0.04] | -0.031 | -2.40 | 0.0166 | 0.0866 |
| L superior parietal | 0.063 | 0.092 | [-0.12,0.24] | 0.007 | 0.69 | 0.4928 | 0.6780 |

| | | | | | | | |
|---|---|---|---|---|---|---|---|
| R superior parietal | 0.013 | 0.092 | [-0.17,0.19] | 0.001 | 0.14 | 0.8917 | 0.9378 |
| L superior temporal | -0.183 | 0.092 | [-0.36,-0.002] | -0.027 | -1.99 | 0.0471 | 0.1780 |
| R superior temporal | -0.235 | 0.092 | [-0.42,-0.05] | -0.034 | -2.55 | 0.0111 | 0.0688 |
| L supramarginal | -0.213 | 0.092 | [-0.4,-0.03] | -0.027 | -2.32 | 0.0209 | 0.1013 |
| R supramarginal | -0.120 | 0.092 | [-0.3,0.06] | -0.015 | -1.30 | 0.1949 | 0.4017 |
| L frontal pole | -0.225 | 0.092 | [-0.41,-0.04] | -0.056 | -2.45 | 0.0147 | 0.0832 |
| R frontal pole | 0.034 | 0.092 | [-0.15,0.22] | 0.007 | 0.37 | 0.7132 | 0.8083 |
| L temporal pole | 0.004 | 0.092 | [-0.18,0.19] | 0.001 | 0.05 | 0.9625 | 0.9625 |
| R temporal pole | -0.038 | 0.092 | [-0.22,0.14] | -0.012 | -0.42 | 0.6778 | 0.8070 |
| L transverse temporal | -0.162 | 0.092 | [-0.34,0.02] | -0.032 | -1.76 | 0.0793 | 0.2396 |
| R transverse temporal | -0.242 | 0.092 | [-0.42,-0.06] | -0.048 | -2.62 | 0.0089 | 0.0688 |
| L insula | -0.104 | 0.092 | [-0.29,0.08] | -0.016 | -1.13 | 0.2587 | 0.4510 |
| R insula | -0.055 | 0.092 | [-0.24,0.13] | -0.009 | -0.60 | 0.5504 | 0.7198 |

**Note.** General linear models per region controlling for age, sex, scan version, and T1w quality (MRIQC efc). FDR corrected using Benjamini-Hochberg procedure across all CT regions. Regions with $p_{FDR}$<0.05 are depicted in bold. S1 = subtype 1, HC = healthy control, SE = standard error, L = left, R = right.

**Table S8b. Cortical thickness comparisons among data-driven subtype 2 and healthy controls.**

| Region | Cohen's *d* (S2 v. HC) | SE | 95% CI | Estimate | *t* value | *p*-value | FDR *p*-value |
|---|---|---|---|---|---|---|---|
| **L banks STS** | **-0.633** | **0.106** | **[-0.84,-0.42]** | **-0.093** | **-6.03** | **2.778e-09** | **2.1004e-08** |
| **R banks STS** | **-0.590** | **0.106** | **[-0.8,-0.38]** | **-0.090** | **-5.63** | **2.765e-08** | **1.4463e-07** |
| **L caudal anterior cingulate** | **-0.293** | **0.105** | **[-0.5,-0.09]** | **-0.053** | **-2.79** | **5.450e-03** | **7.2665e-03** |
| **R caudal anterior cingulate** | **-0.230** | **0.105** | **[-0.44,-0.02]** | **-0.038** | **-2.20** | **0.0285** | **0.0346** |
| **L caudal middle frontal** | **-0.462** | **0.106** | **[-0.67,-0.25]** | **-0.062** | **-4.41** | **1.237e-05** | **2.5496e-05** |
| **R caudal middle frontal** | **-0.513** | **0.106** | **[-0.72,-0.31]** | **-0.067** | **-4.89** | **1.301e-06** | **4.0217e-06** |
| **L cuneus** | **-0.241** | **0.105** | **[-0.45,-0.03]** | **-0.033** | **-2.29** | **0.0221** | **0.0274** |
| **R cuneus** | **-0.273** | **0.105** | **[-0.48,-0.07]** | **-0.033** | **-2.60** | **9.424e-03** | **0.0121** |
| L entorhinal | -0.126 | 0.105 | [-0.33,0.08] | -0.039 | -1.20 | 0.2311 | 0.2455 |
| R entorhinal | -0.174 | 0.105 | [-0.38,0.03] | -0.056 | -1.66 | 0.0967 | 0.1079 |
| **L fusiform** | **-0.667** | **0.107** | **[-0.88,-0.46]** | **-0.077** | **-6.36** | **3.984e-10** | **9.0667e-09** |
| **R fusiform** | **-0.519** | **0.106** | **[-0.73,-0.31]** | **-0.070** | **-4.95** | **9.681e-07** | **3.2917e-06** |
| **L inferior parietal** | **-0.499** | **0.106** | **[-0.71,-0.29]** | **-0.061** | **-4.76** | **2.404e-06** | **6.5402e-06** |
| **R inferior parietal** | **-0.596** | **0.106** | **[-0.81,-0.39]** | **-0.070** | **-5.69** | **2.017e-08** | **1.1430e-07** |
| **L inferior temporal** | **-0.298** | **0.105** | **[-0.5,-0.09]** | **-0.038** | **-2.84** | **4.677e-03** | **6.3605e-03** |
| **R inferior temporal** | **-0.424** | **0.106** | **[-0.63,-0.22]** | **-0.060** | **-4.04** | **6.026e-05** | **1.2053e-04** |
| L isthmus cingulate | -0.196 | 0.105 | [-0.4,0.01] | -0.031 | -1.87 | 0.0622 | 0.0717 |
| R isthmus cingulate | -0.179 | 0.105 | [-0.38,0.03] | -0.028 | -1.70 | 0.0888 | 0.1006 |
| **L lateral occipital** | **-0.633** | **0.106** | **[-0.84,-0.42]** | **-0.068** | **-6.04** | **2.675e-09** | **2.1004e-08** |
| **R lateral occipital** | **-0.639** | **0.106** | **[-0.85,-0.43]** | **-0.070** | **-6.09** | **1.978e-09** | **1.9234e-08** |
| **L lateral orbitofrontal** | **-0.278** | **0.105** | **[-0.48,-0.07]** | **-0.040** | **-2.65** | **8.300e-03** | **0.0109** |
| **R lateral orbitofrontal** | **-0.339** | **0.105** | **[-0.55,-0.13]** | **-0.040** | **-3.24** | **1.274e-03** | **1.9696e-03** |
| **L lingual** | **-0.388** | **0.106** | **[-0.6,-0.18]** | **-0.045** | **-3.70** | **2.339e-04** | **4.0781e-04** |
| **R lingual** | **-0.370** | **0.105** | **[-0.58,-0.16]** | **-0.041** | **-3.53** | **4.461e-04** | **7.5308e-04** |
| L medial orbitofrontal | -0.120 | 0.105 | [-0.33,0.09] | -0.015 | -1.15 | 0.2518 | 0.2635 |
| R medial orbitofrontal | -0.146 | 0.105 | [-0.35,0.06] | -0.017 | -1.40 | 0.1633 | 0.1787 |
| **L middle temporal** | **-0.517** | **0.106** | **[-0.73,-0.31]** | **-0.071** | **-4.93** | **1.036e-06** | **3.3555e-06** |
| **R middle temporal** | **-0.585** | **0.106** | **[-0.79,-0.38]** | **-0.079** | **-5.57** | **3.727e-08** | **1.8103e-07** |
| **L parahippocampal** | **-0.402** | **0.106** | **[-0.61,-0.2]** | **-0.100** | **-3.84** | **1.382e-04** | **2.4734e-04** |
| **R parahippocampal** | **-0.304** | **0.105** | **[-0.51,-0.1]** | **-0.063** | **-2.90** | **3.902e-03** | **5.5280e-03** |
| **L paracentral** | **-0.527** | **0.106** | **[-0.74,-0.32]** | **-0.073** | **-5.02** | **6.712e-07** | **2.4023e-06** |
| **R paracentral** | **-0.408** | **0.106** | **[-0.62,-0.2]** | **-0.057** | **-3.89** | **1.120e-04** | **2.0588e-04** |
| **L pars opercularis** | **-0.554** | **0.106** | **[-0.76,-0.35]** | **-0.082** | **-5.28** | **1.810e-07** | **7.6934e-07** |
| **R pars opercularis** | **-0.529** | **0.106** | **[-0.74,-0.32]** | **-0.079** | **-5.05** | **5.965e-07** | **2.2536e-06** |
| **L pars orbitalis** | **-0.244** | **0.105** | **[-0.45,-0.04]** | **-0.045** | **-2.32** | **0.0205** | **0.0258** |
| **R pars orbitalis** | **-0.310** | **0.105** | **[-0.52,-0.1]** | **-0.058** | **-2.96** | **3.210e-03** | **4.7162e-03** |
| **L pars triangularis** | **-0.414** | **0.106** | **[-0.62,-0.21]** | **-0.062** | **-3.95** | **8.773e-05** | **1.6571e-04** |
| **R pars triangularis** | **-0.501** | **0.106** | **[-0.71,-0.29]** | **-0.077** | **-4.78** | **2.194e-06** | **6.2171e-06** |
| L pericalcarine | -0.028 | 0.105 | [-0.23,0.18] | -0.004 | -0.26 | 0.7931 | 0.7931 |
| R pericalcarine | -0.207 | 0.105 | [-0.41,-0.0005] | -0.027 | -1.97 | 0.0491 | 0.0576 |
| **L postcentral** | **-0.550** | **0.106** | **[-0.76,-0.34]** | **-0.059** | **-5.24** | **2.188e-07** | **8.7532e-07** |
| **R postcentral** | **-0.655** | **0.107** | **[-0.87,-0.45]** | **-0.070** | **-6.25** | **7.678e-10** | **1.0608e-08** |
| **L posterior cingulate** | **-0.370** | **0.105** | **[-0.58,-0.16]** | **-0.049** | **-3.53** | **4.541e-04** | **7.5308e-04** |
| **R posterior cingulate** | **-0.310** | **0.105** | **[-0.52,-0.1]** | **-0.038** | **-2.95** | **3.260e-03** | **4.7162e-03** |
| **L precentral** | **-0.655** | **0.107** | **[-0.86,-0.45]** | **-0.084** | **-6.25** | **7.799e-10** | **1.0608e-08** |
| **R precentral** | **-0.688** | **0.107** | **[-0.9,-0.48]** | **-0.086** | **-6.56** | **1.113e-10** | **5.7800e-09** |
| **L precuneus** | **-0.498** | **0.106** | **[-0.71,-0.29]** | **-0.058** | **-4.75** | **2.587e-06** | **6.7656e-06** |
| **R precuneus** | **-0.464** | **0.106** | **[-0.67,-0.26]** | **-0.055** | **-4.42** | **1.159e-05** | **2.4631e-05** |
| **L rostral anterior cingulate** | **-0.300** | **0.105** | **[-0.51,-0.09]** | **-0.053** | **-2.86** | **4.384e-03** | **6.0836e-03** |
| R rostral anterior cingulate | 0.103 | 0.105 | [-0.1,0.31] | 0.019 | 0.99 | 0.3243 | 0.3342 |
| **L rostral middle frontal** | **-0.418** | **0.106** | **[-0.63,-0.21]** | **-0.049** | **-3.98** | **7.568e-05** | **1.4704e-04** |
| **R rostral middle frontal** | **-0.464** | **0.106** | **[-0.67,-0.26]** | **-0.056** | **-4.43** | **1.121e-05** | **2.4586e-05** |
| **L superior frontal** | **-0.482** | **0.106** | **[-0.69,-0.27]** | **-0.065** | **-4.59** | **5.265e-06** | **1.2786e-05** |
| **R superior frontal** | **-0.505** | **0.106** | **[-0.71,-0.3]** | **-0.071** | **-4.82** | **1.811e-06** | **5.3541e-06** |
| **L superior parietal** | **-0.479** | **0.106** | **[-0.69,-0.27]** | **-0.056** | **-4.57** | **5.901e-06** | **1.3377e-05** |

| | | | | | | | |
|---|---|---|---|---|---|---|---|
| **R superior parietal** | **-0.489** | **0.106** | **[-0.7,-0.28]** | **-0.056** | **-4.66** | **3.806e-06** | **9.5843e-06** |
| **L superior temporal** | **-0.604** | **0.106** | **[-0.81,-0.4]** | **-0.091** | **-5.76** | **1.318e-08** | **8.1476e-08** |
| **R superior temporal** | **-0.682** | **0.107** | **[-0.89,-0.47]** | **-0.100** | **-6.50** | **1.673e-10** | **5.7800e-09** |
| **L supramarginal** | **-0.572** | **0.106** | **[-0.78,-0.36]** | **-0.073** | **-5.45** | **7.143e-08** | **3.2382e-07** |
| **R supramarginal** | **-0.640** | **0.106** | **[-0.85,-0.43]** | **-0.079** | **-6.10** | **1.886e-09** | **1.9234e-08** |
| **L frontal pole** | **-0.354** | **0.105** | **[-0.56,-0.15]** | **-0.089** | **-3.37** | **7.917e-04** | **1.2819e-03** |
| R frontal pole | -0.081 | 0.105 | [-0.29,0.13] | -0.017 | -0.78 | 0.4378 | 0.4443 |
| L temporal pole | -0.208 | 0.105 | [-0.41,-0.002] | -0.064 | -1.99 | 0.0475 | 0.0567 |
| R temporal pole | -0.146 | 0.105 | [-0.35,0.06] | -0.047 | -1.39 | 0.1655 | 0.1787 |
| **L transverse temporal** | **-0.480** | **0.106** | **[-0.69,-0.27]** | **-0.094** | **-4.58** | **5.700e-06** | **1.3366e-05** |
| **R transverse temporal** | **-0.617** | **0.106** | **[-0.83,-0.41]** | **-0.123** | **-5.88** | **6.750e-09** | **4.5900e-08** |
| **L insula** | **-0.346** | **0.105** | **[-0.55,-0.14]** | **-0.054** | **-3.30** | **1.011e-03** | **1.5989e-03** |
| **R insula** | **-0.327** | **0.105** | **[-0.53,-0.12]** | **-0.051** | **-3.12** | **1.921e-03** | **2.9025e-03** |

**Note.** General linear models per region controlling for age, sex, scan version, and T1w quality (MRIQC efc). FDR corrected using Benjamini-Hochberg procedure across all CT regions. Regions with $p_{FDR}<0.05$ are depicted in bold. S2 = subtype 2, HC = healthy control, SE = standard error, L = left, R = right.

**Table S9a. Surface area comparisons among data-driven subtype 1 and healthy controls.**

| Region | Cohen's *d* (S1 v. HC) | SE | 95% CI | Estimate | *t* value | *p*-value | FDR *p*-value |
|---|---|---|---|---|---|---|---|
| L banks STS | 0.122 | 0.092 | [-0.06,0.3] | 19.70 | 1.32 | 0.1866 | 0.1923 |
| **R banks STS** | **0.209** | **0.092** | **[0.03,0.39]** | **26.14** | **2.27** | **0.0236** | **0.0282** |
| **L caudal anterior cingulate** | **0.207** | **0.092** | **[0.03,0.39]** | **26.84** | **2.25** | **0.0249** | **0.0292** |
| R caudal anterior cingulate | 0.152 | 0.092 | [-0.03,0.33] | 22.30 | 1.65 | 0.0990 | 0.1068 |
| **L caudal middle frontal** | **0.257** | **0.092** | **[0.08,0.44]** | **86.56** | **2.79** | **5.4882e-03** | **7.6163e-03** |
| **R caudal middle frontal** | **0.271** | **0.092** | **[0.09,0.45]** | **92.09** | **2.94** | **3.3679e-03** | **5.2050e-03** |
| **L cuneus** | **0.364** | **0.093** | **[0.18,0.55]** | **72.48** | **3.95** | **8.8491e-05** | **3.3430e-04** |
| **R cuneus** | **0.423** | **0.093** | **[0.24,0.61]** | **83.53** | **4.59** | **5.3017e-06** | **3.6054e-05** |
| L entorhinal | 0.099 | 0.092 | [-0.08,0.28] | 8.82 | 1.07 | 0.2856 | 0.2898 |
| R entorhinal | 0.122 | 0.092 | [-0.06,0.3] | 9.64 | 1.33 | 0.1847 | 0.1923 |
| **L fusiform** | **0.295** | **0.093** | **[0.11,0.48]** | **97.90** | **3.20** | **1.4270e-03** | **2.6227e-03** |
| **R fusiform** | **0.297** | **0.093** | **[0.12,0.48]** | **86.89** | **3.23** | **1.3223e-03** | **2.5690e-03** |
| **L inferior parietal** | **0.229** | **0.092** | **[0.05,0.41]** | **128.78** | **2.49** | **0.0132** | **0.0170** |
| R inferior parietal | 0.185 | 0.092 | [0.004,0.37] | 123.34 | 2.01 | 0.0452 | 0.0513 |
| **L inferior temporal** | **0.233** | **0.092** | **[0.05,0.42]** | **101.11** | **2.53** | **0.0116** | **0.0151** |
| **R inferior temporal** | **0.346** | **0.093** | **[0.16,0.53]** | **131.99** | **3.76** | **1.8754e-04** | **5.9737e-04** |
| **L isthmus cingulate** | **0.414** | **0.093** | **[0.23,0.6]** | **58.97** | **4.49** | **8.5431e-06** | **5.2811e-05** |
| **R isthmus cingulate** | **0.354** | **0.093** | **[0.17,0.54]** | **44.13** | **3.84** | **1.3427e-04** | **4.6193e-04** |
| **L lateral occipital** | **0.382** | **0.093** | **[0.2,0.56]** | **214.50** | **4.14** | **3.9285e-05** | **1.8137e-04** |
| **R lateral occipital** | **0.285** | **0.093** | **[0.1,0.47]** | **162.56** | **3.10** | **2.0455e-03** | **3.4773e-03** |
| **L lateral orbitofrontal** | **0.378** | **0.093** | **[0.2,0.56]** | **107.00** | **4.10** | **4.6249e-05** | **1.9656e-04** |
| **R lateral orbitofrontal** | **0.275** | **0.093** | **[0.09,0.46]** | **88.01** | **2.99** | **2.9353e-03** | **4.7523e-03** |
| **L lingual** | **0.264** | **0.092** | **[0.08,0.45]** | **99.60** | **2.87** | **4.2581e-03** | **6.1017e-03** |
| **R lingual** | **0.194** | **0.092** | **[0.01,0.38]** | **77.03** | **2.10** | **0.0359** | **0.0413** |
| **L medial orbitofrontal** | **0.405** | **0.093** | **[0.22,0.59]** | **80.62** | **4.40** | **1.2941e-05** | **7.3332e-05** |
| **R medial orbitofrontal** | **0.433** | **0.093** | **[0.25,0.62]** | **86.99** | **4.70** | **3.1559e-06** | **3.6054e-05** |
| **L middle temporal** | **0.215** | **0.092** | **[0.03,0.4]** | **80.55** | **2.33** | **0.0201** | **0.0248** |
| **R middle temporal** | **0.300** | **0.093** | **[0.12,0.48]** | **119.93** | **3.26** | **1.1809e-03** | **2.3618e-03** |
| L parahippocampal | 0.170 | 0.092 | [-0.01,0.35] | 11.89 | 1.85 | 0.0651 | 0.0725 |
| **R parahippocampal** | **0.245** | **0.092** | **[0.06,0.43]** | **18.06** | **2.66** | **8.0737e-03** | **0.0108** |
| **L paracentral** | **0.296** | **0.093** | **[0.12,0.48]** | **45.51** | **3.22** | **1.3669e-03** | **2.5819e-03** |
| **R paracentral** | **0.323** | **0.093** | **[0.14,0.51]** | **54.05** | **3.51** | **4.7885e-04** | **1.2524e-03** |
| **L pars opercularis** | **0.301** | **0.093** | **[0.12,0.48]** | **73.88** | **3.27** | **1.1460e-03** | **2.3615e-03** |
| **R pars opercularis** | **0.276** | **0.093** | **[0.09,0.46]** | **52.63** | **2.99** | **2.8670e-03** | **4.7523e-03** |
| **L pars orbitalis** | **0.310** | **0.093** | **[0.13,0.49]** | **27.13** | **3.37** | **7.9711e-04** | **1.7485e-03** |
| **R pars orbitalis** | **0.319** | **0.093** | **[0.14,0.5]** | **33.75** | **3.46** | **5.6907e-04** | **1.4332e-03** |
| **L pars triangularis** | **0.250** | **0.092** | **[0.07,0.43]** | **47.28** | **2.72** | **6.7942e-03** | **9.2401e-03** |
| **R pars triangularis** | **0.270** | **0.092** | **[0.09,0.45]** | **59.37** | **2.94** | **3.4489e-03** | **5.2117e-03** |
| **L pericalcarine** | **0.314** | **0.093** | **[0.13,0.5]** | **77.16** | **3.41** | **6.9244e-04** | **1.6237e-03** |
| **R pericalcarine** | **0.264** | **0.092** | **[0.08,0.45]** | **65.73** | **2.87** | **4.3071e-03** | **6.1017e-03** |
| **L postcentral** | **0.345** | **0.093** | **[0.16,0.53]** | **144.58** | **3.75** | **1.9327e-04** | **5.9737e-04** |
| **R postcentral** | **0.433** | **0.093** | **[0.25,0.62]** | **172.38** | **4.70** | **3.1466e-06** | **3.6054e-05** |
| **L posterior cingulate** | **0.216** | **0.092** | **[0.03,0.4]** | **33.63** | **2.34** | **0.0196** | **0.0247** |
| **R posterior cingulate** | **0.373** | **0.093** | **[0.19,0.56]** | **58.02** | **4.04** | **5.9112e-05** | **2.3645e-04** |
| **L precentral** | **0.332** | **0.093** | **[0.15,0.51]** | **158.06** | **3.60** | **3.4223e-04** | **9.6965e-04** |
| **R precentral** | **0.423** | **0.093** | **[0.24,0.61]** | **181.11** | **4.60** | **5.2297e-06** | **3.6054e-05** |
| **L precuneus** | **0.516** | **0.093** | **[0.33,0.7]** | **213.71** | **5.60** | **3.2319e-08** | **2.1760e-06** |
| **R precuneus** | **0.492** | **0.093** | **[0.31,0.68]** | **212.49** | **5.34** | **1.3202e-07** | **4.4880e-06** |
| **L rostral anterior cingulate** | **0.307** | **0.093** | **[0.13,0.49]** | **47.62** | **3.34** | **9.0119e-04** | **1.9150e-03** |
| R rostral anterior cingulate | 0.083 | 0.092 | [-0.1,0.26] | 10.10 | 0.90 | 0.3689 | 0.3689 |
| **L rostral middle frontal** | **0.427** | **0.093** | **[0.24,0.61]** | **270.07** | **4.63** | **4.4134e-06** | **3.6054e-05** |
| **R rostral middle frontal** | **0.425** | **0.093** | **[0.24,0.61]** | **294.32** | **4.62** | **4.7817e-06** | **3.6054e-05** |
| **L superior frontal** | **0.381** | **0.093** | **[0.2,0.56]** | **289.68** | **4.14** | **4.0007e-05** | **1.8137e-04** |
| **R superior frontal** | **0.354** | **0.093** | **[0.17,0.54]** | **265.48** | **3.84** | **1.3586e-04** | **4.6193e-04** |
| **L superior parietal** | **0.452** | **0.093** | **[0.27,0.64]** | **281.31** | **4.91** | **1.1771e-06** | **2.6679e-05** |

| | | | | | | | |
|---|---|---|---|---|---|---|---|
| **R superior parietal** | **0.386** | **0.093** | **[0.2,0.57]** | **222.31** | **4.19** | **3.2532e-05** | **1.7017e-04** |
| **L superior temporal** | **0.332** | **0.093** | **[0.15,0.51]** | **138.96** | **3.61** | **3.3400e-04** | **9.6965e-04** |
| **R superior temporal** | **0.314** | **0.093** | **[0.13,0.5]** | **114.41** | **3.41** | **6.8243e-04** | **1.6237e-03** |
| **L supramarginal** | **0.326** | **0.093** | **[0.14,0.51]** | **196.61** | **3.54** | **4.3549e-04** | **1.1845e-03** |
| **R supramarginal** | **0.432** | **0.093** | **[0.25,0.61]** | **200.25** | **4.69** | **3.3997e-06** | **3.6054e-05** |
| **L frontal pole** | **0.311** | **0.093** | **[0.13,0.49]** | **10.53** | **3.37** | **7.8961e-04** | **1.7485e-03** |
| **R frontal pole** | **0.265** | **0.092** | **[0.08,0.45]** | **11.47** | **2.88** | **4.1046e-03** | **6.0676e-03** |
| **L temporal pole** | **0.211** | **0.092** | **[0.03,0.39]** | **14.09** | **2.29** | **0.0225** | **0.0273** |
| R temporal pole | 0.159 | 0.092 | [-0.02,0.34] | 10.83 | 1.73 | 0.0845 | 0.0927 |
| **L transverse temporal** | **0.288** | **0.093** | **[0.11,0.47]** | **17.97** | **3.13** | **1.8204e-03** | **3.1740e-03** |
| **R transverse temporal** | **0.291** | **0.093** | **[0.11,0.47]** | **12.73** | **3.16** | **1.6596e-03** | **2.9698e-03** |
| L insula | 0.125 | 0.092 | [-0.06,0.31] | 34.15 | 1.36 | 0.1741 | 0.1849 |
| **R insula** | **0.272** | **0.092** | **[0.09,0.45]** | **59.40** | **2.96** | **3.2239e-03** | **5.0983e-03** |

**Note.** General linear models per region controlling for age, sex, scan version, and T1w quality (MRIQC efc). FDR corrected using Benjamini-Hochberg procedure across all SA regions. Regions with $p_{FDR}$<0.05 are depicted in bold. S1 = subtype 1, HC = healthy control, SE = standard error, L = left, R = right.

**<u>Table S9b.</u> Surface area comparisons among data-driven subtype 2 and healthy controls.**

| Region | Cohen's *d* (S2 v. HC) | SE | 95% CI | Estimate | *t* value | *p*-value | FDR *p*-value |
|---|---|---|---|---|---|---|---|
| **L banks STS** | **-0.476** | **0.106** | **[-0.68,-0.27]** | **-77.04** | **-4.54** | **6.7392e-06** | **8.6465e-06** |
| **R banks STS** | **-0.491** | **0.106** | **[-0.7,-0.28]** | **-61.39** | **-4.68** | **3.5292e-06** | **4.7997e-06** |
| **L caudal anterior cingulate** | **-0.422** | **0.106** | **[-0.63,-0.22]** | **-54.74** | **-4.03** | **6.3264e-05** | **7.0524e-05** |
| **R caudal anterior cingulate** | **-0.485** | **0.106** | **[-0.69,-0.28]** | **-71.09** | **-4.63** | **4.5178e-06** | **6.0238e-06** |
| **L caudal middle frontal** | **-0.577** | **0.106** | **[-0.79,-0.37]** | **-194.49** | **-5.50** | **5.5622e-08** | **9.0055e-08** |
| **R caudal middle frontal** | **-0.479** | **0.106** | **[-0.69,-0.27]** | **-162.75** | **-4.57** | **5.9107e-06** | **7.7293e-06** |
| **L cuneus** | **-0.615** | **0.106** | **[-0.82,-0.41]** | **-122.55** | **-5.86** | **7.4979e-09** | **1.3925e-08** |
| **R cuneus** | **-0.531** | **0.106** | **[-0.74,-0.32]** | **-104.74** | **-5.06** | **5.5616e-07** | **8.0466e-07** |
| **L entorhinal** | **-0.425** | **0.106** | **[-0.63,-0.22]** | **-38.07** | **-4.05** | **5.7027e-05** | **6.4631e-05** |
| **R entorhinal** | **-0.502** | **0.106** | **[-0.71,-0.29]** | **-39.58** | **-4.79** | **2.1096e-06** | **2.9886e-06** |
| **L fusiform** | **-0.787** | **0.107** | **[-1,-0.58]** | **-261.05** | **-7.50** | **2.1747e-13** | **1.3444e-12** |
| **R fusiform** | **-1.003** | **0.109** | **[-1.22,-0.79]** | **-293.40** | **-9.57** | **2.5895e-20** | **1.7680e-18** |
| **L inferior parietal** | **-0.691** | **0.107** | **[-0.9,-0.48]** | **-388.78** | **-6.59** | **9.4184e-11** | **2.5618e-10** |
| **R inferior parietal** | **-0.726** | **0.107** | **[-0.94,-0.52]** | **-484.52** | **-6.92** | **1.1059e-11** | **3.5810e-11** |
| **L inferior temporal** | **-0.902** | **0.108** | **[-1.11,-0.69]** | **-390.86** | **-8.60** | **6.5763e-17** | **8.9438e-16** |
| **R inferior temporal** | **-0.919** | **0.108** | **[-1.13,-0.71]** | **-350.55** | **-8.77** | **1.7820e-17** | **4.0392e-16** |
| **L isthmus cingulate** | **-0.604** | **0.106** | **[-0.81,-0.4]** | **-86.07** | **-5.76** | **1.3597e-08** | **2.3116e-08** |
| **R isthmus cingulate** | **-0.548** | **0.106** | **[-0.76,-0.34]** | **-68.35** | **-5.23** | **2.3512e-07** | **3.7182e-07** |
| **L lateral occipital** | **-0.613** | **0.106** | **[-0.82,-0.4]** | **-344.43** | **-5.84** | **8.3563e-09** | **1.4953e-08** |
| **R lateral occipital** | **-0.747** | **0.107** | **[-0.96,-0.54]** | **-425.79** | **-7.13** | **2.9215e-12** | **1.1686e-11** |
| **L lateral orbitofrontal** | **-0.714** | **0.107** | **[-0.92,-0.51]** | **-202.25** | **-6.81** | **2.2880e-11** | **7.0721e-11** |
| **R lateral orbitofrontal** | **-0.697** | **0.107** | **[-0.91,-0.49]** | **-223.06** | **-6.65** | **6.5200e-11** | **1.9276e-10** |
| **L lingual** | **-0.629** | **0.106** | **[-0.84,-0.42]** | **-236.96** | **-6.00** | **3.4445e-09** | **7.0977e-09** |
| **R lingual** | **-0.543** | **0.106** | **[-0.75,-0.34]** | **-215.91** | **-5.18** | **3.0394e-07** | **4.6973e-07** |
| **L medial orbitofrontal** | **-0.829** | **0.108** | **[-1.04,-0.62]** | **-164.93** | **-7.90** | **1.2734e-14** | **1.0824e-13** |
| **R medial orbitofrontal** | **-0.651** | **0.107** | **[-0.86,-0.44]** | **-130.75** | **-6.21** | **9.7128e-10** | **2.2016e-09** |
| **L middle temporal** | **-0.930** | **0.108** | **[-1.14,-0.72]** | **-349.07** | **-8.87** | **7.7755e-18** | **2.6438e-16** |
| **R middle temporal** | **-0.907** | **0.108** | **[-1.12,-0.7]** | **-362.47** | **-8.65** | **4.4270e-17** | **7.5259e-16** |
| **L parahippocampal** | **-0.418** | **0.106** | **[-0.63,-0.21]** | **-29.20** | **-3.99** | **7.5162e-05** | **8.2435e-05** |
| **R parahippocampal** | **-0.461** | **0.106** | **[-0.67,-0.25]** | **-34.00** | **-4.39** | **1.3062e-05** | **1.5861e-05** |
| **L paracentral** | **-0.605** | **0.106** | **[-0.81,-0.4]** | **-92.97** | **-5.77** | **1.2482e-08** | **2.1763e-08** |
| **R paracentral** | **-0.686** | **0.107** | **[-0.9,-0.48]** | **-114.68** | **-6.54** | **1.2577e-10** | **3.2735e-10** |
| **L pars opercularis** | **-0.353** | **0.105** | **[-0.56,-0.15]** | **-86.67** | **-3.37** | **8.0785e-04** | **8.1991e-04** |
| **R pars opercularis** | **-0.534** | **0.106** | **[-0.74,-0.33]** | **-101.96** | **-5.09** | **4.6615e-07** | **7.0440e-07** |
| **L pars orbitalis** | **-0.762** | **0.107** | **[-0.97,-0.55]** | **-66.58** | **-7.27** | **1.1295e-12** | **5.4861e-12** |
| **R pars orbitalis** | **-0.591** | **0.106** | **[-0.8,-0.38]** | **-62.55** | **-5.64** | **2.5947e-08** | **4.3034e-08** |
| **L pars triangularis** | **-0.393** | **0.106** | **[-0.6,-0.19]** | **-74.24** | **-3.75** | **1.9609e-04** | **2.0515e-04** |
| **R pars triangularis** | **-0.463** | **0.106** | **[-0.67,-0.26]** | **-101.61** | **-4.41** | **1.1993e-05** | **1.4827e-05** |
| **L pericalcarine** | **-0.460** | **0.106** | **[-0.67,-0.25]** | **-113.05** | **-4.39** | **1.3436e-05** | **1.6029e-05** |
| **R pericalcarine** | **-0.411** | **0.106** | **[-0.62,-0.2]** | **-102.43** | **-3.92** | **9.7627e-05** | **1.0373e-04** |
| **L postcentral** | **-0.783** | **0.107** | **[-0.99,-0.57]** | **-327.53** | **-7.46** | **2.9022e-13** | **1.6446e-12** |
| **R postcentral** | **-0.778** | **0.107** | **[-0.99,-0.57]** | **-309.34** | **-7.42** | **4.0204e-13** | **2.1030e-12** |
| **L posterior cingulate** | **-0.642** | **0.107** | **[-0.85,-0.43]** | **-100.27** | **-6.13** | **1.6078e-09** | **3.4165e-09** |
| **R posterior cingulate** | **-0.686** | **0.107** | **[-0.9,-0.48]** | **-106.79** | **-6.54** | **1.2998e-10** | **3.2735e-10** |
| **L precentral** | **-0.739** | **0.107** | **[-0.95,-0.53]** | **-352.05** | **-7.05** | **4.9562e-12** | **1.8724e-11** |
| **R precentral** | **-0.807** | **0.107** | **[-1.02,-0.6]** | **-345.41** | **-7.70** | **5.4592e-14** | **3.7123e-13** |
| **L precuneus** | **-0.833** | **0.108** | **[-1.04,-0.62]** | **-345.17** | **-7.95** | **9.2665e-15** | **9.0017e-14** |
| **R precuneus** | **-0.866** | **0.108** | **[-1.08,-0.66]** | **-374.23** | **-8.26** | **8.9995e-16** | **1.0199e-14** |
| **L rostral anterior cingulate** | **-0.625** | **0.106** | **[-0.83,-0.42]** | **-96.82** | **-5.96** | **4.3127e-09** | **8.6254e-09** |
| **R rostral anterior cingulate** | **-0.756** | **0.107** | **[-0.97,-0.55]** | **-92.20** | **-7.21** | **1.6764e-12** | **7.1246e-12** |
| **L rostral middle frontal** | **-0.727** | **0.107** | **[-0.94,-0.52]** | **-460.11** | **-6.93** | **1.0462e-11** | **3.5569e-11** |
| **R rostral middle frontal** | **-0.680** | **0.107** | **[-0.89,-0.47]** | **-470.59** | **-6.48** | **1.8550e-10** | **4.5050e-10** |
| **L superior frontal** | **-0.732** | **0.107** | **[-0.94,-0.52]** | **-556.23** | **-6.98** | **7.7326e-12** | **2.7675e-11** |
| **R superior frontal** | **-0.818** | **0.107** | **[-1.03,-0.61]** | **-614.07** | **-7.80** | **2.6235e-14** | **1.9822e-13** |
| **L superior parietal** | **-0.758** | **0.107** | **[-0.97,-0.55]** | **-471.71** | **-7.23** | **1.4385e-12** | **6.5214e-12** |

| | | | | | | | |
|---|---|---|---|---|---|---|---|
| **R superior parietal** | **-0.692** | **0.107** | **[-0.9,-0.48]** | **-399.07** | **-6.60** | **8.8312e-11** | **2.5022e-10** |
| **L superior temporal** | **-0.644** | **0.107** | **[-0.85,-0.44]** | **-269.43** | **-6.14** | **1.4409e-09** | **3.1607e-09** |
| **R superior temporal** | **-0.666** | **0.107** | **[-0.88,-0.46]** | **-242.47** | **-6.36** | **4.0480e-10** | **9.4918e-10** |
| **L supramarginal** | **-0.614** | **0.106** | **[-0.82,-0.41]** | **-370.79** | **-5.86** | **7.5770e-09** | **1.3925e-08** |
| **R supramarginal** | **-0.533** | **0.106** | **[-0.74,-0.33]** | **-247.22** | **-5.08** | **4.9108e-07** | **7.2594e-07** |
| **L frontal pole** | **-0.412** | **0.106** | **[-0.62,-0.2]** | **-13.95** | **-3.92** | **9.6560e-05** | **1.0373e-04** |
| **R frontal pole** | **-0.469** | **0.106** | **[-0.68,-0.26]** | **-20.29** | **-4.48** | **9.0568e-06** | **1.1405e-05** |
| **L temporal pole** | **-0.329** | **0.105** | **[-0.54,-0.12]** | **-21.98** | **-3.13** | **1.8021e-03** | **1.8021e-03** |
| **R temporal pole** | **-0.376** | **0.105** | **[-0.58,-0.17]** | **-25.56** | **-3.58** | **3.6960e-04** | **3.8080e-04** |
| **L transverse temporal** | **-0.501** | **0.106** | **[-0.71,-0.29]** | **-31.24** | **-4.78** | **2.1747e-06** | **3.0179e-06** |
| **R transverse temporal** | **-0.426** | **0.106** | **[-0.63,-0.22]** | **-18.65** | **-4.06** | **5.4434e-05** | **6.2738e-05** |
| **L insula** | **-0.452** | **0.106** | **[-0.66,-0.24]** | **-123.04** | **-4.31** | **1.9268e-05** | **2.2590e-05** |
| **R insula** | **-0.619** | **0.106** | **[-0.83,-0.41]** | **-135.00** | **-5.90** | **5.8888e-09** | **1.1441e-08** |

**Note.** General linear models per region controlling for age, sex, scan version, and T1w quality (MRIQC efc). FDR corrected using Benjamini-Hochberg procedure across all SA regions. Regions with $p_{FDR}<0.05$ are depicted in bold. S2 = subtype 2, HC = healthy control, SE = standard error, L = left, R = right.

**Table S10a.** Subcortical volume comparisons among data-driven subtype 1 and healthy controls.

| Region | Cohen's *d* (S1 v. HC) | SE | 95% CI | Estimate | *t* value | *p*-value | FDR *p*-value |
|---|---|---|---|---|---|---|---|
| L thalamus | 0.003 | 0.094 | [-0.18,0.19] | 1.64 | 0.03 | 0.97 | 0.98 |
| R thalamus | -0.02 | 0.094 | [-0.21,0.16] | -10.53 | -0.22 | 0.82 | 0.96 |
| **L caudate** | **0.49** | **0.095** | **[0.3,0.67]** | **186.63** | **5.17** | **3.12e-07** | **4.43e-06** |
| **R caudate** | **0.44** | **0.095** | **[0.25,0.63]** | **172.60** | **4.69** | **3.42e-06** | **2.39e-05** |
| **L putamen** | **0.24** | **0.094** | **[0.05,0.43]** | **141.29** | **2.55** | **0.01** | **0.03** |
| **R putamen** | **0.4** | **0.095** | **[0.21,0.58]** | **175.98** | **4.24** | **2.53e-05** | **1.18e-04** |
| **L pallidum** | **0.25** | **0.094** | **[0.06,0.43]** | **51.68** | **2.61** | **0.01** | **0.03** |
| **R pallidum** | **0.25** | **0.094** | **[0.06,0.43]** | **45.91** | **2.62** | **0.01** | **0.03** |
| L hippocampus | -0.002 | 0.094 | [-0.19,0.18] | -0.76 | -0.03 | 0.98 | 0.98 |
| R hippocampus | 0.06 | 0.094 | [-0.13,0.24] | 17.97 | 0.61 | 0.54 | 0.82 |
| L amygdala | 0.13 | 0.094 | [-0.06,0.31] | 24.95 | 1.36 | 0.17 | 0.31 |
| R amygdala | -0.027 | 0.094 | [-0.21,0.16] | -5.49 | -0.29 | 0.78 | 0.96 |
| L accumbens | 0.05 | 0.094 | [-0.13,0.24] | 5.16 | 0.55 | 0.59 | 0.82 |
| R accumbens | 0.15 | 0.094 | [-0.04,0.33] | 12.87 | 1.59 | 0.11 | 0.23 |

**Note.** General linear models per region controlling for age, sex, scan version, T1w quality (MRIQC efc), and intracranial volume. *$p$<0.05, **$p$<0.01, ***$p$<0.001. FDR corrected using Benjamini-Hochberg procedure across all SV regions. Regions with $p_{FDR}$<0.05 are depicted in bold. S1 = subtype 1, HC = healthy control, SE = standard error, L = left, R = right.

**Table S10b.** Subcortical volume comparisons among data-driven subtype 2 and healthy controls.

| Region | Cohen's *d* (S2 v. HC) | SE | 95% CI | Estimate | *t* value | *p*-value | FDR *p*-value |
|---|---|---|---|---|---|---|---|
| **L thalamus** | **-0.79** | **0.112** | **[-1.01,-0.57]** | **-433.46** | **-7.22** | **1.51e-12** | **4.21e-12** |
| **R thalamus** | **-0.77** | **0.112** | **[-0.99,-0.55]** | **-387.45** | **-7.02** | **5.98e-12** | **1.40e-11** |
| **L caudate** | **-0.33** | **0.110** | **[-0.55,-0.11]** | **-126.30** | **-3.00** | **0.003** | **0.004** |
| **R caudate** | **-0.34** | **0.110** | **[-0.55,-0.12]** | **-131.60** | **-3.06** | **0.002** | **0.003** |
| **L putamen** | **-0.33** | **0.110** | **[-0.55,-0.11]** | **-194.80** | **-3.01** | **0.003** | **0.004** |
| R putamen | -0.17 | 0.110 | [-0.38,0.05] | -72.98 | -1.51 | 0.13 | 0.13 |
| L pallidum | -0.19 | 0.110 | [-0.4,0.03] | -39.14 | -1.70 | 0.09 | 0.10 |
| R pallidum | -0.21 | 0.110 | [-0.43,0.01] | -39.17 | -1.91 | 0.06 | 0.07 |
| **L hippocampus** | **-0.93** | **0.113** | **[-1.15,-0.7]** | **-290.32** | **-8.44** | **2.34e-16** | **1.61e-15** |
| **R hippocampus** | **-0.93** | **0.113** | **[-1.15,-0.71]** | **-291.35** | **-8.47** | **1.82e-16** | **1.61e-15** |
| **L amygdala** | **-0.89** | **0.113** | **[-1.11,-0.67]** | **-173.39** | **-8.08** | **3.41e-15** | **1.59e-14** |
| **R amygdala** | **-0.83** | **0.112** | **[-1.05,-0.61]** | **-170.29** | **-7.57** | **1.41e-13** | **4.94e-13** |
| **L accumbens** | **-0.53** | **0.111** | **[-0.75,-0.32]** | **-53.74** | **-4.86** | **1.50e-06** | **2.62e-06** |
| **R accumbens** | **-0.55** | **0.111** | **[-0.77,-0.33]** | **-47.61** | **-5.03** | **6.46e-07** | **1.29e-06** |

**Note.** General linear models per region controlling for age, sex, scan version, T1w quality (MRIQC efc), and intracranial volume. *$p$<0.05, **$p$<0.01, ***$p$<0.001. FDR corrected using Benjamini-Hochberg procedure across all SV regions. Regions with $p_{FDR}$<0.05 are depicted in bold. S2 = subtype 2, HC = healthy control, SE = standard error, L = left, R = right.

REFERENCES

1. First, M., Spitzer, R., Gibbon, M., Williams, J., *Structured clinical interview for DSM-IV-TR axis I disorders, research version, patient edition (SCID-I/P)*. New York: Biometrics Research, 2002.
